# Tracing the origins of human voltage-gated K^+^ channels: where they come from and what we lost along the way

**DOI:** 10.1101/2025.09.21.677649

**Authors:** Zhaoyang Jiang, Neha Kopri, Alla Zurnachyan, Timothy Jegla

## Abstract

Humans have 40 voltage-gated K^+^ channel (Kv) genes that are broadly conserved in vertebrate model organisms and spread across three ancient gene families (KCNQ, EAG and Shaker). We used deep coverage phylogenetic analyses to trace the evolutionary origins of each human Kv channel. 39/40 human channels were already present in the common ancestor of jawed vertebrates (gnathostomes) and as many as 36 could have an origin in a genome duplication within the stem gnathostome lineage. The Elk channel Kv12.2 is the newest human Kv stemming from a gene duplication in the ancestor of tetrapods and lobe-finned fish. Kv5.1, is the oldest human Kv channel, traceable to the stem chordate lineage. We found evidence for a total of 45 widespread gnathostome Kv channels. Humans lost five channels that can still be found in some other gnathostome lineages. Birds and mammals have the smallest remaining Kv channel sets among gnathostomes and a lineage-specific genome duplication in teleost fish increased their ancestral Kv set to as many as 69 channels. We expressed orthologs of the 5 gnathostome channels missing in humans from vampire bat (Kv3.5), harpy eagle (Kv7.1L), duck-billed platypus (Kv8.3), coelacanth (Kv9.4) and axolotl (Kv12.4). Each has some unique functional characteristics, but their gating properties are largely redundant with existing human Kv paralogs. The human Kv set still contains substantial gating redundancy and we argue that loss-of-function mutations driving partitioning and subfunctionalization have probably had a greater influence on the broad retention of ancestral gnathostome Kv channels than gain-of-function gating optimizations.

**SUMMARY STATEMENT:** We used deep coverage phylogenies to trace the origins of 36/40 human voltage-gated K^+^ channel genes to a common ancestor of jawed vertebrates. We identified and functionally characterized 5 additional channels that humans lost from other living vertebrates.

## INTRODUCTION

Voltage-gated K^+^ channels (Kv channels) are essential for neuronal excitability and many other aspects of our physiology. Human Kv channels display a broad range of gating phenotypes, subcellular localizations and cell type-specific expression patterns that diversify electrical signaling patterns our nervous systems can generate. The 40 human genes that encode pore-forming subunits of Kv channels are distributed across three gene families: Shaker (Kv1-6,8,9), KCNQ (Kv7) and EAG (Kv10-12) (Jegla et al., 2009). The Shaker family comprises the majority of the human Kv channel set at 27 genes, while KCNQ and EAG have 5 and 8 genes, respectively. We will use broadly accepted IUPHAR nomenclature (Kv1.x-12.x) (Gutman et al., 2003) when referring to individual human Kv channels and their orthologs, but it is important to remember this nomenclature does not reflect all evolutionary relationships. The Shaker, KCNQ and EAG families derive from distinct molecular lineages that last had common ancestors in microbes (Jegla et al., 2018; Jegla et al., 2024; Simonson et al., 2025) and they do not form a single clade to the exclusion of other voltage-gated ion channels in phylogenies (Yu and Catterall, 2004; Jegla et al., 2018).

Subunits from all three families do share a common core motif consisting of a voltage sensor domain (VSD) couple to a pore domain (PD), and they function as tetramers with a single composite pore surrounded by four independent VSDs (Long et al., 2005; Whicher and MacKinnon, 2016; Sun and MacKinnon, 2017). However, they can easily be differentiated in sequence searches by 1) amino acid conservation level within this common core and 2) presence of family-specific cytoplasmic domains that regulate gating and assembly (Lara et al., 2023).

Shaker family channels have an N-terminal T1 domain that regulates assembly and subunit compatibility (Xu et al., 1995; Kreusch et al., 1998; Nanao et al., 2003), KCNQs have a C-terminal coiled-coil domain that serves a similar function (Jenke et al., 2003; Schwake et al., 2006; Sun and MacKinnon, 2017), and EAG channels have an N-terminal PAS domain (eag domain) and a C-terminal C-linker/cyclic nucleotide binding homology domain (CNBHD) that regulate gating (Morais Cabral et al., 1998; Brelidze et al., 2012). The eag domain has been lost independently in a subset of cnidarian and bilaterian Erg (Kv11) channels (Martinson et al., 2014), but these are still easily identifiable in sequence searches.

The human Shaker and EAG gene families can be further subdivided into the Kv1-4 (Shaker, Shab, Shaw and Shal) and the Kv10-12 (Eag, Erg and Elk) gene subfamilies, respectively. These subfamilies form separate clades in molecular phylogenies that are traceable to evolutionary origins in the common ancestor of cnidarians and bilaterians (Jegla et al., 1995; Jegla and Salkoff, 1997; Sand et al., 2011; Jegla et al., 2012; Martinson et al., 2014; Li et al., 2015b; Li et al., 2015c; Lara et al., 2023). One of the most functionally significant attributes of these gene subfamilies is that they are assembly exclusive; there is no mixing of subunits across subfamilies boundaries during assembly (Covarrubias et al., 1991; Wimmers et al., 2001; Zou et al., 2003).

Heteromeric assembly does however occur within subfamilies and the vertebrate Kv2 subfamily even includes an expansion of “silent” subunits (Kv5, Kv6, Kv8 and Kv9 channels) that require co-assembly with Kv2.1 and Kv2.2 α-subunits to form functional channels (Post et al., 1996; Jegla et al., 2012; Li et al., 2015b; Bocksteins, 2016; Pisupati et al., 2018). Subfamily restricted assembly allows multiple independent Shaker and EAG family channel types to be expressed in a single cell without interaction and thus allows more complex signal patterning. In humans, the KCNQ family has two assembly groups that could also be seen as representing distinct subfamilies: Kv7.2-7.5 can form heteromultimers, but they do no co-assemble with Kv7.1 (Wang et al., 1998; Wickenden et al., 2001; Schwake et al., 2006; Howard et al., 2007). There is phylogenetic evidence for both putative KCNQ subfamilies in bilaterian invertebrates (Jegla et al., 2009; Li et al., 2015b), but the cnidarian/bilaterian ancestor only had a single type of KCNQ channel and ctenophores, the basal animal lineage (Ryan et al., 2013; Schultz et al., 2023), lack KCNQ channels entirely (Li et al., 2015b; Lara et al., 2023; Simonson et al., 2025). This split of the KCNQ family is remarkably the only known bilaterian-specific Kv channel subfamily addition despite the evolution of complex centralized nervous systems specifically within Bilateria. Humans, and probably most bilaterians, can theoretically express up to 9 independent Kv channel types in a single cell based on these subfamily assembly restrictions. The common ancestor of cnidarians and bilaterians, which is believed to have used a comparatively simple diffuse nerve net (Rolls and Jegla, 2015; Stone et al., 2020; Stone et al., 2025), already had 8 of these subfamilies (Lara et al., 2023; Simonson et al., 2025). Ctenophores in contrast lack our Kv subfamilies, but there are indications that they might have evolved their own subfamilies independently within a large ctenophore-restricted expansion of the Shaker family (Li et al., 2015b; Simonson et al., 2024; Simonson et al., 2025).

Individual human Kv genes are much newer than the subfamilies to which they belong and are shared only with other vertebrates. For example, the voltage-gated K^+^ channel set of humans and mice are identical (Jegla et al., 2009), making mouse the key model system for genetic analysis of Kv physiology. In contrast, bilaterian invertebrates typically have fewer Kv channels and thus lack 1:1 gene orthology with vertebrates (Jegla et al., 2009). This distribution pattern suggests that most of our voltage-gated K^+^ channels could be derived from two whole genome duplications that occurred in the stem vertebrate lineage (Ohno, 1970, 1999; Panopoulou et al., 2003; Dehal and Boore, 2005). The genes produced by these duplications are often referred to as “ohnologs”, in recognition of Susumu Ohno who first proposed these duplications as the “2R” hypothesis. The phylogenetic spread of a large expansion of genes in the Shaker Kv2 subfamily is indeed consistent with a major role for the vertebrate genome duplications (Jegla and Simonson, 2023), but this has not been confirmed with phylogenetics. Tracing the origins of vertebrate genes to Ohno’s genome duplications isn’t as simple as counting genes because extensive gene loss typically occurs after genome duplications (Wolfe, 2001; Makino and McLysaght, 2012). Thus the predicted 4:1 (AB)(CD) pattern in vertebrate gene phylogenies expected for a simple 2R genome duplication scenario is uncommon (Friedman and Hughes, 2001). Deviations from the (AB)(CD) pattern can also be generated by unequal selection pressures, or subsequent duplications of individual genes, as documented for voltage-gated Na^+^ channels (Zakon et al., 2011).

These complications have presented a barrier to tracing the duplication history and origins of each human Kv channel gene, but this barrier can be overcome or at least minimized with broad species coverage in molecular phylogenies. Increasing species coverage is particularly important for identifying gene losses since any one draft genome may have missing data. This deep species coverage in phylogenies is now possible with the recent availability of multiple high-quality genomes and transcriptomes from every major vertebrate lineage as well as most major bilaterian invertebrate lineages. Thus, we can now determine the timing of the gene duplications that produced the human Kv channel set relative to the divergence order of bilaterian lineages.

Furthermore, analysis of genomes from the earliest vertebrate lineage, cyclostomes or jawless vertebrates such as lampreys and hagfish, have also provided new insights into the two timing of Ohno’s genome duplications (Smith et al., 2013; Marlétaz et al., 2024; Yu et al., 2024). The first duplication (1R) occurred in a vertebrate common ancestor prior to the divergence of cyclostomes from gnathostomes (jawed vertebrates). In contrast, the second duplication (2R) occurred in a common ancestor of gnathostomes (Marlétaz et al., 2024; Yu et al., 2024) after divergence from cyclostomes. Genes duplications that occurred after vertebrates separated from tunicates but prior to the divergence of cyclostomes are candidates for a 1R origin, whereas duplications in the gnathostome common ancestor are candidates for a 2R origin. A third genome duplication (3R) occurred in the teleost lineage of ray-finned fish (Taylor et al., 2001; Taylor et al., 2003; Jaillon et al., 2004) and is the reason why zebrafish often have co-orthologs for human genes.

Here we built Kv channel phylogenies with deep species coverage of vertebrates and bilaterian invertebrates and find that most human Kvs are gnathostome-specific and likely to be 2R ohnologs. In contrast, the contribution of the 1R genome duplication and additional duplications of individual genes was much more limited. Most interestingly, we found 5 additional ancient vertebrate Kv channels humans have lost at various points in our evolutionary history. The vertebrate lineage leading to humans thus at one point had 45 Kv genes, though no living vertebrates have retained all 45 and losses are most extensive in mammals and birds.

Functional expression of orthologs of the 5 Kv channels humans are largely redundant, though not identical to, existing human paralogs at the level of gating biophysics. Given the similar level of gating redundancy remaining in human Kvs, we argue that other factors like functional partitioning and subfunctionalization must have played a significant role in keeping the human Kv channel set as large as it is.

## METHODS

### Sequence collection

All Kv sequences collected for phylogenetic analysis are provided in plain text FASTA format as Data S1-S6. Complete Kv channel sets for some bilaterians including humans, mouse and Drosophila are well-established (Wei et al., 1996; Gutman et al., 2003; Jegla et al., 2009) and served as our query sequences in BLAST searches (Altschul et al., 1997) for other bilaterian species. Previously identified Kv channels from cnidarians and ctenophores (Jegla et al., 2012; Li et al., 2015b; Li et al., 2015c; Lara et al., 2023; Simonson et al., 2025) were used as outgroups. Proteins predicted from animal genomes are usually accurate and complete, so we were able to collect full Kv sets from most target species with BLASTP searches of the NCBI NR database where these predictions are deposited (Sayers et al., 2021). We used relaxed BLAST search parameters (Expect threshold = 1 and a word size = 3) which we have found reliably detect all related gene family members (Baker et al., 2015; Li et al., 2015b; Jegla et al., 2016; Lara et al., 2023; Jegla et al., 2024; Simonson et al., 2025). Supplemental TBLASTN sequence searches of the source genome drafts were used to ensure all Kv loci had been identified, and rarely proteins from additional loci without gene predictions were manually assembled using sequence homology and intron/exon boundary predictions as guides. A single human protein query sequence from each of the three Kv gene families is sufficient to find all members of the gene family, but we often used multiple query sequences from each gene subfamily to pre-sort hits for phylogenetic analysis. It was occasionally necessary to perform a reciprocal BLAST search with identified sequences as queries against mouse or human REFSEQ (O’Leary et al., 2016) to confirm subfamily identity. There are a wide variety of search methods available for comprehensive bulk sequence collection, but we find serial BLAST searches with carefully selected query sequences provide a highly efficient workflow and yield the most accurate and data-rich results.

For incomplete protein predictions, we filled gaps in the conserved regions used for phylogenetic analysis with data from genome drafts or transcriptomes whenever possible. If remaining gaps were > 10% of the region used for phylogenetic analysis, the sequence was discarded. In most cases we were able to substitute discarded sequences with orthologs from an alternate species in the same genus or family and we used multiple species to represent key animal clades whenever possible. None of the lineage-specific gene losses cited in our analysis can be explained by discarded sequences.

### Phylogenetic analysis

We used amino acid sequences for phylogenetic analysis because they have the best resolving power over the long evolutionary time frames. The trade-off is that ion channel orthologs in closely related species like birds or placental mammals sometimes do not have enough informative sequence positions to resolve branch order. However, this did not compromise our analysis because there are no recent Kv gene duplications in the tetrapod lineage. Main amino acid alignments used for phylogenetic analysis were made in Mega12 (Kumar et al., 2024) with Muscle (Edgar, 2004) adjusted to a gap open penalty of ≥ −5, and manually trimmed to remove un-conserved regions with length variation or spurious alignment. For extended length vertebrate-only phylogenies, alignments were made in MAFFT (Katoh and Standley, 2013) with a gap open penalty of 7 and local pairing, and then trimmed with Gblocks (Castresana, 2000) with a 50% coverage threshold for conserved positions and a requirement of ≥ 3 consecutive conserved residues for inclusion in the final alignment. Both strategies work well on conserved core domains with few gaps, but the MAFFT/Gblocks pipeline is more efficient and accurate for maximal extension of alignments into regions where blocks of conservation are more widely dispersed.

All phylogenies were run by two independent methods: Bayesian inference (BI) as implemented using Mr. Bayes v3.2.7 (Ronquist et al., 2012) or maximum likelihood (ML) implemented using IQ-TREE 2 (Minh et al., 2020). Phylogenetic analysis in Mr. Bayes was GPU-accelerated using Beagle 3 (Ayres et al., 2019) and run under a mixed amino acid model to allow selection of the best model for the data sets. The first 25 percent of the analysis was discarded as burn-in and at this point all runs had flat likelihood and total branch length trajectories. We used two runs of 6 chains for BI phylogenies and ran them for 3,000,000 generations, sampling the statistics every 5,000 generations. The Kv1 subfamily phylogeny took longer to converge and ran for a total of 6,000,000 generations. Alignment lengths, taxa numbers, run convergence parameters and amino acid models are given in Table S1. Node support in the BI phylogenies is expressed as posterior probability (PP). For maximum likelihood, we used the model finder function in IQ-TREE 2 to select the best fit model for each analysis (Table S1). Node support was tested with 10,000 ultrafast bootstrap (BS) replications.

### Cloning and RNA preparation

ORFs for harpy eagle Kv7.1L (Hharp_Kv7.1L), axolotl Kv12.4 (Amexi_Kv12.4), platypus Kv8.3 (Oanat_Kv8.3), coelacanth Kv9.4 (Lchal_Kv9.4) and vampire bat Kv3.1 and Kv3.5 (Drotu_Kv3.1/3.5) were synthesized (Genscript, Piscataway, NJ) using Xenopus-optimized codons, bracketed with Xenopus β-globin UTRs from the pOX expression vector (Jegla and Salkoff, 1997), cloned into pUC57 and sequence verified. Oocyte expression vectors for human Kv7.2 (Li et al., 2015b), Kv12.1-12.3 (Zhang et al., 2009; Kazmierczak et al., 2013), and Kv2.1 (Pisupati et al., 2018) have been previously described. Oocyte expression vectors for Kv8.2 and Kv9.3 encode proteins identical to corresponding sequences in the Data S4 FASTA file and were made over 20 years ago from Origene (Rockville, MD) cDNA clones identified based on homology to mouse brain transcripts (Gustincich et al., 2003), but are being used for the first time here. Templates for *in vitro* transcription were generated by high-fidelity PCR using a **F**orward primer to the 5′ UTR incorporating a T7 promoter (underlined) modified for incorporation of CleanCap-AG (TriLink Biotechnologies, San Diego, CA) and a **R**everse primer to the 3′ UTR that adds a short poly-A tail:

**F:** 5′-CCGCGAAATTAATACGACTCACTATAAGTTGTTCTTTTTGCAGAAGCTCAGAAT-3′

**R:** 5′-TTTTTTTTTTTTTTTTTTTTTTTTTTTTTTGTGAAGAAACTTTCTTTTTATTAGGAGCAG-3′

For run-off *in vitro* transcription, 0.4–1 μg purified templates were transcribed with 100 U T7 polymerase, 20 U RNase inhibitor (Takara, San Jose, CA or ThermoFisher, Waltham, MA), 4 mM CleanCap-AG, 5 mM DTT (Sigma-Aldrich, St. Louis, MO), a 20 mM balanced nucleotide mix, and 0.02 U yeast inorganic pyrophosphatase (NEB, Ipswich, MA) for 1–2 hours at 37°C. Transcripts were purified by LiCl precipitation, resuspended in RNase inhibitor-supplemented water (1:20 dilution), and analyzed by gel electrophoresis to verify transcript integrity. RNA yields typically ranged from 20–60 μg per reaction. In our hands, this protocol produces higher protein expression/ng transcribed RNA than commercially available *in vitro* transcription kits (Simonson et al., 2025).

### Oocytes and Electrophysiology

Mature oocytes were isolated from intact *Xenopus laevis* ovaries (Xenopus 1, Ann Arbor, MI) with 0.5–1 mg/mL type II collagenase in calcium-free ND98 saline (98 mM NaCl, 2 mM KCl, 1 mM MgCl₂, 5 mM HEPES; pH 7.2). Isolated oocytes were maintained in a cool room (∼18°C) in ND98 culture medium (ND98 saline supplemented with 1.8 mM CaCl₂, 2.5 mM Na-pyruvate, 100 U/mL penicillin, 100 µg/mL streptomycin, and 50 µg/mL tetracycline). For functional expression experiments, oocytes were microinjected with 50 nL of RNA solution containing 0.1– 20 ng of each transcript and incubated for 24–72h prior to recording.

Two-electrode voltage-clamp (TEVC) recordings were performed at room temperature (20-23℃) under continuous perfusion (0.5–1 mL/min) of a low-chloride recording bath solution (98 mM NaOH, 2 mM KCl, 1 mM CaCl_2_, and 5 mM HEPES, adjusted to pH 7.2 with methane sulfonic acid). Microelectrodes were pulled from borosilicate glass capillaries and filled with 3 M KCl, giving tip resistances of 0.5-1 MΩ. The bath reference (Ag/AgCl) was coupled to the chamber through a 1M NaCl salt bridge. Signals were collected with a CA-1B amplifier (Dagan Instruments, Minneapolis, MN), digitized via a Digidata 1440A interface, and processed using pClamp 10 software (Molecular Devices, Sunnyvale, CA). Data acquisition was performed at 5– 10 kHz with low-pass filtering at 5 kHz. Series resistance was compensated as needed for larger currents.

Data and statistical analyses were performed using Clampfit 10.3 (Molecular Devices, Sunnyvale, CA) or Origin2025 (OriginLab, Northampton, MA). Conductance–voltage (GV) and steady-state inactivation (SSI) data from individual cells were fitted with a single Boltzmann function (Eq. 1) to derive midpoint (V_50_) and slope (s) values. Data were normalized using the upper (*A1*) and lower (*A2*) asymptotes prior to statistical analyses.

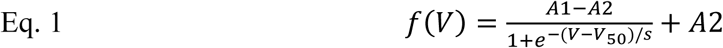

Deactivation time constants were determined using single exponential fits (Eq. 2) of tails currents, where *A* is the amplitude, *b* is the baseline and τ is the time constant. The voltage

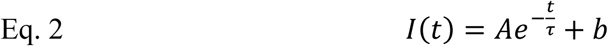

dependence of deactivation was estimated from fits of semi-log plotted τ_deact_ vs. voltage data with the linear form of a single exponential (Eq. 3), where z is the elemental charge associated with deactivation, F is Faraday’s constant, R is the gas constant, T is absolute temperature and τ(0) is the time constant at 0 mV. We derived *z* from the slope of the linear fit (Eq. 4).

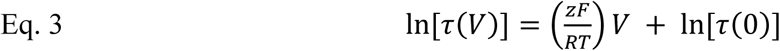

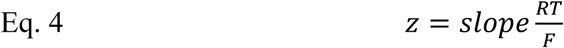

All individual measurements used for electrophysiological analyses are either included in figures or provided in File S1 if figure panels display averages.

## RESULTS

Fig. 1 shows a phylogeny for the main vertebrate species and invertebrate clades we included in our analyses based on the current consensus view of the animal phylogeny with approximate clade divergence times adopted from TimeTree5 (Hedges et al., 2006; Hedges et al., 2015; Kumar et al., 2022). We selected representative species based on 1) genome data quality as assessed in preliminary BLAST searches and 2) author favorites when there multiple high quality options. We used multiple species from each clade of interest whenever possible to deconvolute true gene losses from missing data. Additional species were added to individual phylogenies as needed to fill in missing data or strengthen the evidence for clade-specific gene losses. The main goal of our study was to trace the evolutionary history of human Kv channels within the vertebrates, but we also included broad coverage of protostome invertebrate species to gain preliminary insights into the history of Kv diversification within protostomes. Cnidarian and ctenophore sequences were included as outgroups for tree rooting. These early diverging animals have surprisingly large and diverse Kv channel sets (Jegla et al., 2012; Li et al., 2015b; Li et al., 2015c; Lara et al., 2023; Simonson et al., 2025), so we selected a few representative sequences for computational efficiency. For each phylogeny, we analyzed the alignment with two independent phylogenetic approaches, Bayesian inference (BI) and maximum likelihood (ML) as detailed in the methods to better assess the strength of findings.

**Figure 1.**
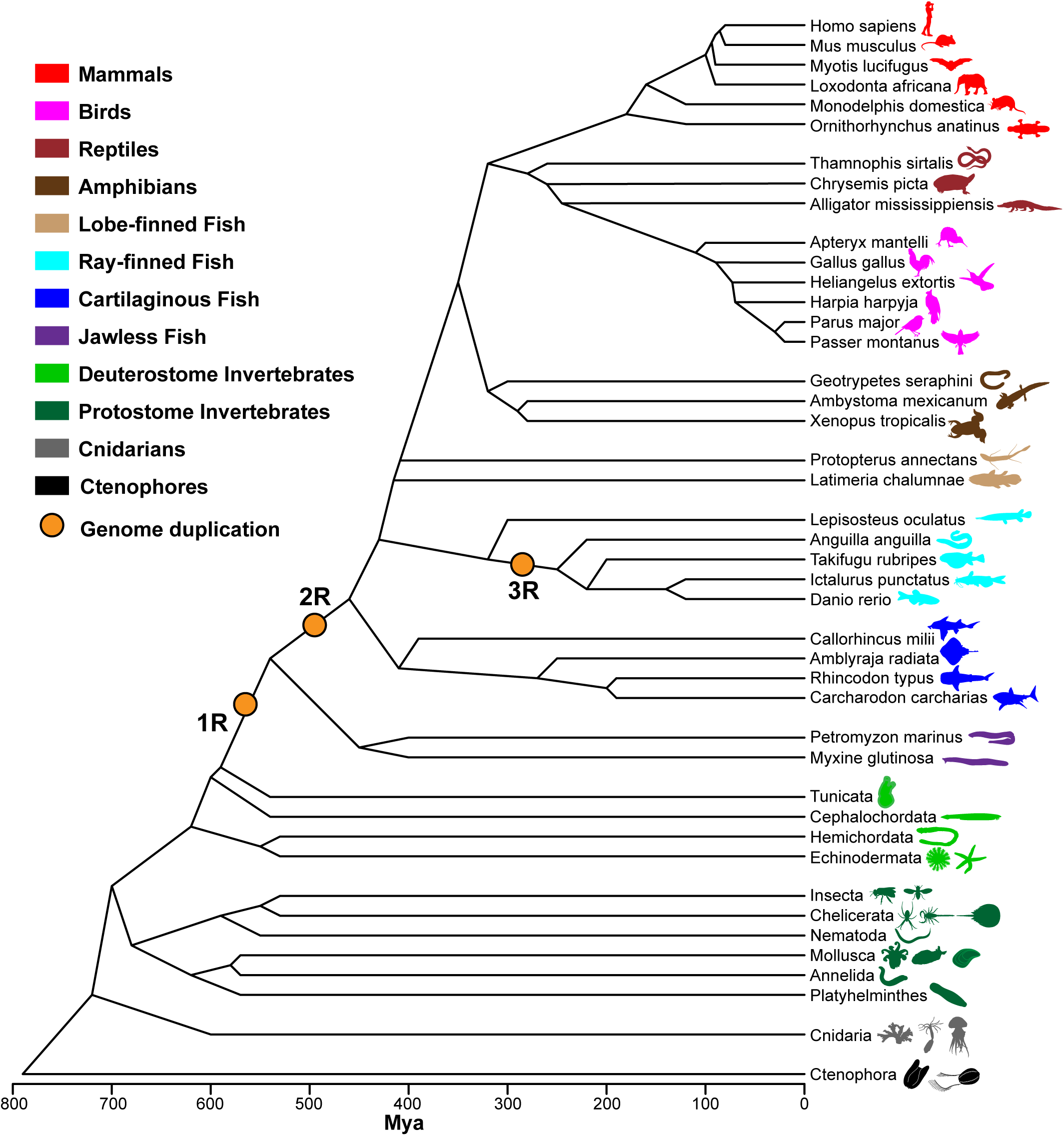
Phylogeny of bilaterian animals. Summary diagram of the current consensus view of evolutionary relationships of bilaterian animals with approximate clade divergence times in millions of years ago (Mya) taken from TimeTree5 estimates (Kumar et al., 2022). The positions of three documented genome duplications in vertebrates (1R-3R) are indicated with orange circles. Key bilaterian clades are colored according to the legend and cnidarians (gray) and ctenophores (black) are included as outgroups. Species names are given for the main vertebrate representatives we used, but note we included alternate and additional species in individual phylogenies as needed. Additional species and all invertebrate species are identified in Data S1-36. Animal icons at the left margin are public domain images sourced from the PhyloPic.org repository.

### KCNQ Family

Fig.2 shows the BI phylogeny of the animal KCNQ gene family with statistical support overlaid at key nodes from both the BI and ML trees. We considered nodes highly supported if they had ≥ 0.95 posterior probability (PP) in BI and ≥ 0.95 ultrafast bootstrap (BS) support in ML. The sequence alignment and original BI and ML tree files with fully expanded clades and complete node support data are provided as Data S7-S9. Separate bilaterian KCNQ1 and KCNQ2 clades are supported in both phylogenies, and we label them as distinct gene subfamilies here because they meet the same functional definition of subfamilies used within the Shaker and EAG gene families: an evolutionarily unified clade of channels that represents a distinct assembly group. Both KCNQ subfamilies have a broad phylogenetic distribution in protostome and deuterostome invertebrates, though most invertebrate species have a single gene for each (Table 1). Nematodes are the notable exception with two KCNQ1 subfamily genes; the highly divergent nematode KQT3 channel (Wei et al., 2005) appears to have arisen from a nematode-specific KCNQ1 family gene duplication. Cnidarian KCNQs fell within the KCNQ2 subfamily rather than forming an outgroup in both phylogenies, so we hypothesize that the KCNQ1 subfamily derives from a bilaterian-specific duplication of an ancestral KCNQ2 subfamily channel.

**Figure 2.**
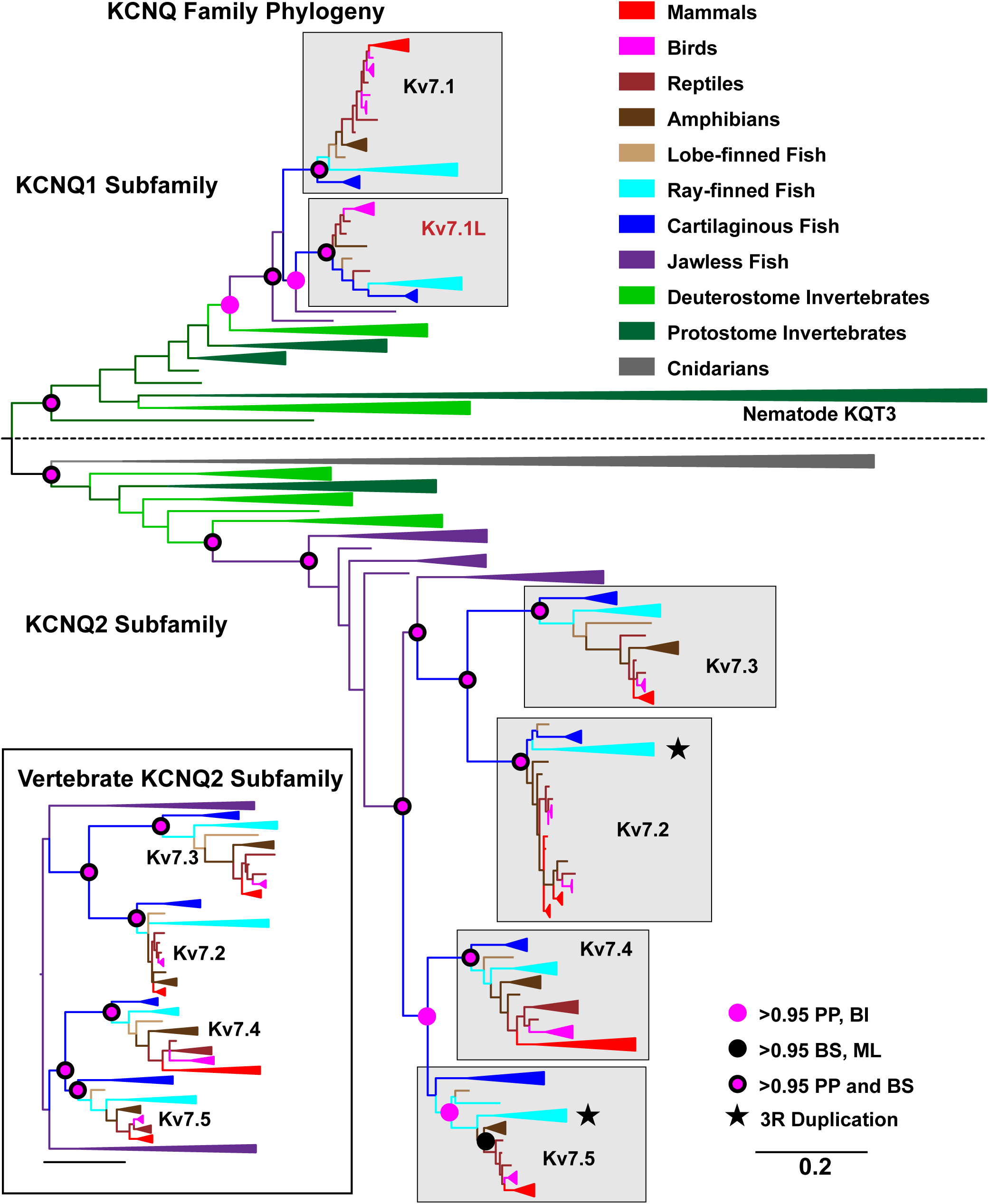
Phylogenetic analysis of the KCNQ gene family. The main figure shows a Bayesian inference phylogeny of the animal KCNQs based on a 398-site alignment of 244 cnidarian and bilaterian sequences. Taxa are colored by phylogenetic group according to the legend and channel clades containing multiple sequences from the same group are collapsed (triangular wedges) for display. Wedge width is not scaled to sequence number. Ortholog groups including human genes are shaded gray and labeled according to the IUPHAR nomenclature and stars mark clades with putative 3R duplications in teleosts. We named a new vertebrate ortholog group missing in mammals Kv7.1L according to its pairing with Kv7.1 in the phylogenies. The KCNQ1 and KCNQ2 gene subfamilies are labeled and separated with a dashed line and the tree is rooted between these subfamilies for display. The nematode-specific KQT3 clade mentioned in the main text is also labeled. Branch lengths reflect substitutions/site according to the scale bar and the length of collapsed wedges is scaled to the longest interior branch. Statistical support of > 0.95 for select key nodes is indicated with colored circles according to the legend. Support is expressed as posterior probability (PP) this Bayesian inference (BI) phylogeny and as bootstrap support (BS) for a maximum likelihood (ML) phylogeny based on the same alignment. (Inset) Bayesian inference phylogeny of 128 vertebrate KCNQ2 subfamily channels based on a 530-site alignment.

**Table 1.**
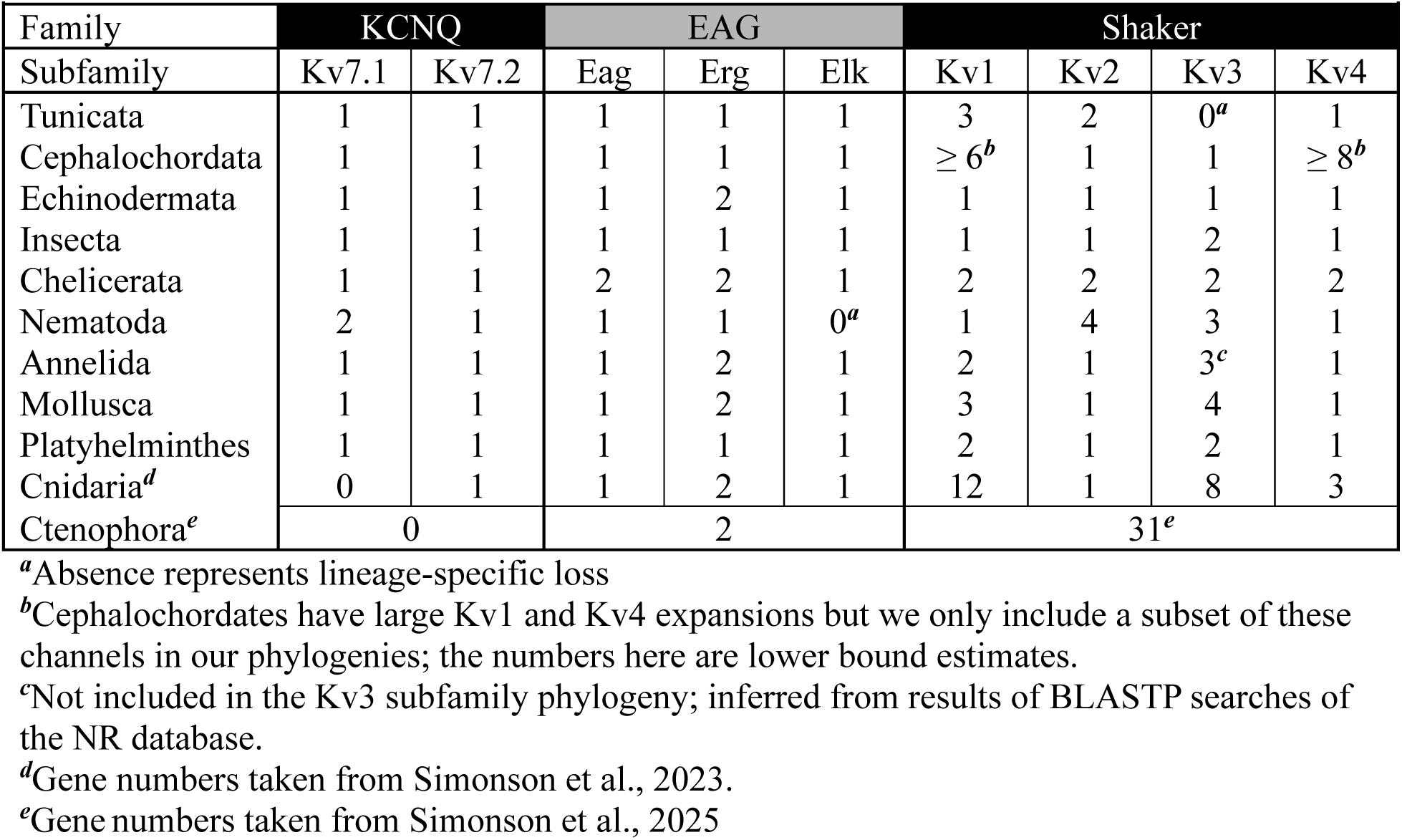
Ancestral Kv channel counts for various invertebrate clades.

The KCNQ1 subfamily which is represented by a single gene (Kv7.1) in mouse and humans unexpectedly contains *two* channels that have a wide distribution in gnathostomes. Kv7.1 is present in all major gnathostome clades, but a second gnathostome channel which we name Kv7.1L here is absent in mammalian genomes. The cyclostomes sea lamprey and atlantic hagfish have a single KCNQ1 subfamily channel suggesting that Kv7.1 and Kv7.1L are 2R ohnologs duplicated from a single ancestral vertebrate channel. We find no evidence to support retention of multiple 1R ohnologs in the KCNQ1 subfamily. There is strong support in both phylogenies for gnathostome-wide Kv7.2-Kv7.4 ortholog clades and for pairing of Kv7.2 and Kv7.3 clades. A unified gnathostome Kv7.5 clade was only present in the Bayesian inference phylogeny, and support for inclusion of cartilaginous fish sequences is weak (PP = 0.64). We reasoned this could be due to the aggressive alignment trimming required for phylogenetic analyses that include both KCNQ subfamilies as well as evolutionarily diverse invertebrate sequences. We therefore ran a second phylogenetic analysis including only vertebrate KCNQ2 subfamily channels which allowed us to increase the alignment length from 398 to 530 positions. This additional alignment and BI and ML trees are provided as Data S10-12. In this second analysis, both BI (Fig. 2, inset) and ML support a unified gnathostome Kv7.5 clade and pairing of Kv7.4 with Kv7.5. In these analyses, Kv7.2-7.5 form the classic (AB)(CD) phylogenetic topology expected for two rounds of genome duplication, and the presence of all four in the stem gnathostome supports a 2R origin. Kv7.2/Kv7.3 and Kv7.4/Kv7.5 clades do not consistently pair with cyclostome sequences at > 0.95 support in either set of phylogenetic analyses. Therefore, we cannot be certain that the Kv7.2-7.5 clade also includes a 1R duplication. An alternative explanation would be loss of one of the two 1R ohnologs coupled with an independent duplication in gnathostomes prior to 2R.

We also looked for retention of 3R KCNQ ohnologs in teleost fish. The alligator gar, *Lepisosteus oculatus*, diverged from teleosts prior to the 3R genome duplication, so 3R duplications can be detected by looking for teleost co-orthologs for single Lepisosteus channels. We defined teleost channels as putative 3R ohnologs if we observed a duplication in at least two of the four teleost species we included in our analyses: *Anguilla anguilla* (eel), *Danio rerio* (zebrafish), *Ictalurus punctatus* (catfish) and *Takifugu rubripes* (pufferfish). We found evidence for 3R ohnologs for Kv7.2 and Kv7.5 (Fig. 2, stars) but not the other four gnathostome Kv7 channels. The teleost common ancestor therefore probably had 8 KCNQ family channels.

We expressed the Kv7.1L ortholog from Harpy eagle (*Harpia harpyja*, Hharp-Kv7.1L1) in *Xenopus* oocytes to see what type of current mammals lost. Birds and reptiles have the closest extant Kv7.1L orthologs to the ancestral Kv7.1L channel that was lost in mammals. We also co-expressed Hharp-Kv7.1L1 with the accessory subunit KCNE1 which co-assembles with human Kv7.1 to generate the classic slowly-activating cardiac I_KS_ current (Barhanin et al., 1996; Sanguinetti et al., 1996). KCNE family subunits modify expression of diverse Kv channels (McDonald et al., 1997; Abbott et al., 1999; McCrossan and Abbott, 2004; Clancy et al., 2009), but the KCNE1-Kv7.1 association is the most well-studied so we wanted to see if it was conserved in Kv7.1L. We used human KCNE1 (Hsapi-KCNE1) for co-expression experiments because its ability to enhance native Xenopus KCNQ1 currents in oocytes suggests there are unlikely to be barriers to cross-species co-assembly (Sanguinetti et al., 1996). Example traces for Kv7.1L, KCNE1 and Kv7.1L + KCNE1 are shown in Fig. 3A. The induction of native KCNQ currents by exogenous KCNE1 was small in our assays, so these native currents do not significantly contaminate our Kv7.1L + KCNE1 currents. The average peak current size at 60 mV for KCNE1 expressed in isolation was just 0.067 ± 0.060 μA (S.D., n = 5) compared to 13.44 ± 5.24 μA (S.D., n = 10) for Kv7.1L + KCNE1. Kv7.1L homomeric currents were slightly smaller (7.51 ± 4.22 μA, n = 7, *p* = 0.03, t-test) and show a classic hook in tail currents that is typically indicative of rapid recovery from inactivation (Fig. 3A, left, arrow). Human Kv7.1 homomers have inactivation that produces a similar tail hook (Tristani-Firouzi and Sanguinetti, 1998). KCNE1 clearly slowed activation of Kv7.1L, but we did not make quantitative comparisons due to the confound of this inactivation. KCNE1 significantly hyperpolarized the V_50_ of Kv7.1L (Fig. 3B, Table 2; *p* < 0.001, t-test), possibly by significantly reducing the charge associated with deactivation (z) (*p* < 0.001, t-test, n = 5) and thereby slowing deactivation at low voltages (Fig. 3C). In contrast, KCNE1 introduces a depolarized shift in human Kv7.1, though KCNE2 and KCNE3 can hyperpolarize Kv7.1 activation (Schroeder et al., 2000; Tinel et al., 2000; Wang et al., 2012; Barro-Soria et al., 2015; Barro-Soria et al., 2017). Confirmation of this opposite GV shift with a species-matched Hharp-KCNE1 would be needed to determine if it represents a real functional difference between Kv7.1 and Kv7.1L, but our results here clearly show the potential KCNE1 regulation is shared between Kv7.1 and Kv7.1L.

**Figure 3.**
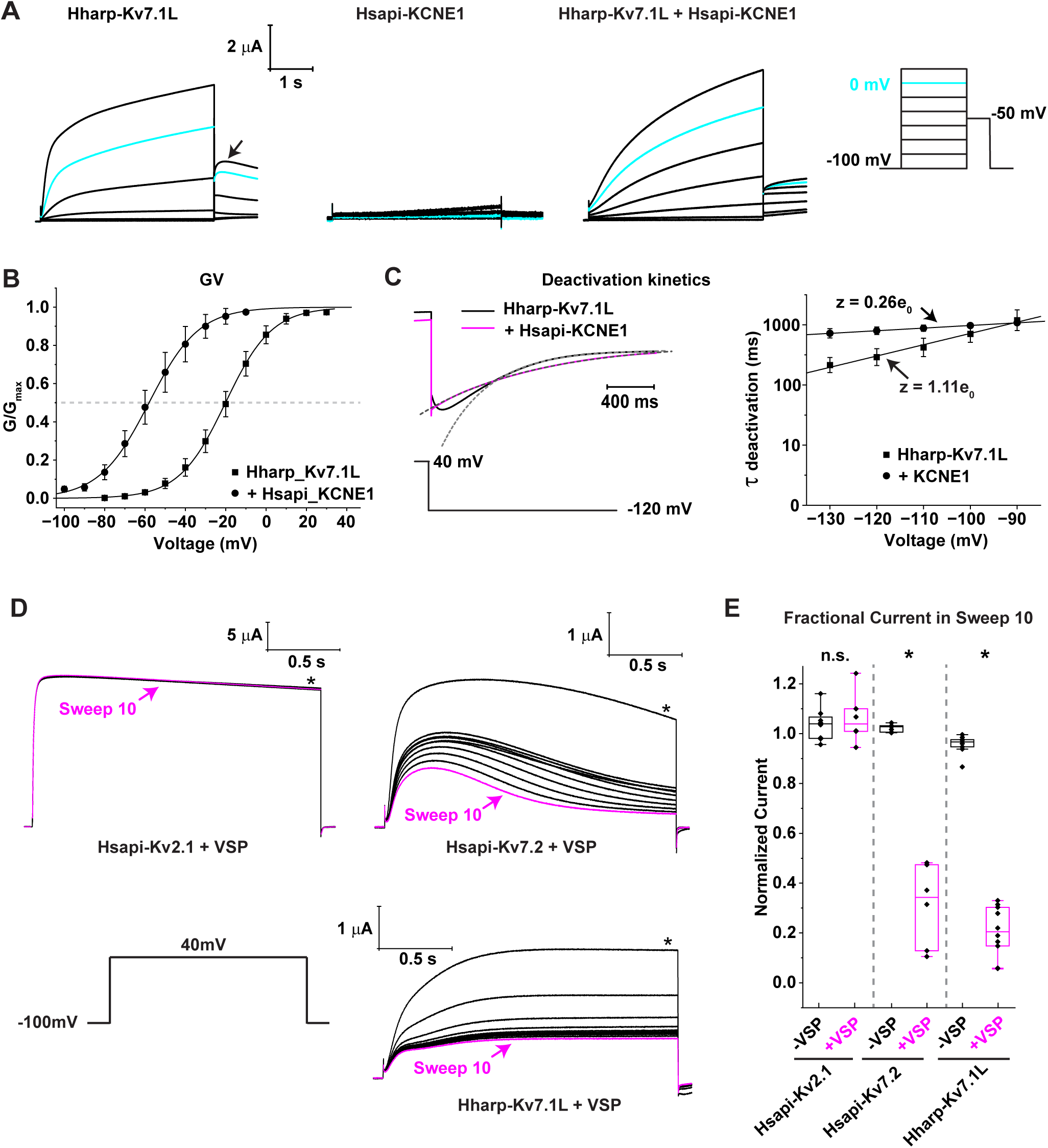
Functional characterization of harpy eagle Kv7.1. (A) Families of currents recorded from oocytes expressing Hharp-Kv7.1L (left), Hsapi-KCNE1 (middle) or both (right) recorded in response to a series of depolarizing voltage steps (protocol cartoon at right margin). The 0 mV trace in each family is highlighted with cyan and the scale bars indicate current amplitude and time. (B) Normalized voltage activation curves for Hharp-Kv7.1L with and without co-expression of Hsapi_KCNE1 measured from isochronal tail currents. Data show mean ± S.D. of 5-6 oocytes. Smooth curves show single Boltzmann fits with the mean V_50_ and slope values from Table 2. (C) Time constants for Hharp-Kv7.1L deactivation were determined with single exponential fits of tail currents recorded at the indicated voltages after a test pulse to +40 mV. The left panel shows examples of fits (dotted gray lines) to normalized tail currents, and the right panel shows a semi-log plot of the deactivation time constant vs. voltage, fit with a single exponential equation to estimate the gating charge (z) associated with deactivation. Data are mean ± S.D., n = 5. (D) Example current traces recorded in response to repeated depolarizing voltage steps to 40 mV in the presence of Ciona VSP reduces Hsapi-Kv7.2 and Hharp-Kv7.1L currents but not Hsapi-Kv2.1 currents. The 10^th^ sweep is highlighted in magenta and asterisks indicate the region of the sweeps used to determine fractional current inhibition. (E) Plots of fractional current size in the 10^th^ sweep compared to the 1^st^ sweep for each channel from oocytes expressing just the channel (black) or the channel + Ciona VSP (magenta). Boxes show median and middle quartiles, individual data points are shown with black diamonds and whiskers mark the observed data range. Statistical significance for current suppression in the presence of VSP is indicated at the top (t-test; n.s., *p* > 0.05; *, *p* < 0.001).

**Table 2.** Boltzmann fit parameters.

| **<u>Conductance-voltage (GV) analyses</u>** |  |  |  |
| --- | --- | --- | --- |
| **Channel** | **V<sub>50</sub> (mV)** | **Slope (mV)** | **n** |
| Hharp-Kv7.1L | -20.1 ± 3.5* | 11.5 ± 0.19 | 6 |
| Hharp-Kv7.1L + Hsapi-KCNE1 | -58.0 ± 4.7 | 12.3 ± 2.2 | 5 |
| Hsapi-Kv12.1 | -48.4 ± 1.7 | 12.0 ± 0.80 | 8 |
| Hsapi-Kv12.2 | -20.4 ± 3.0 | 25.3 ± 2.7 | 6 |
| Hsapi-Kv12.3 | -83.2 ± 1.5 | 13.5 ± 0.90 | 7 |
| Amexi-Kv12.4 | -24.9 ± 5.2 | 14.8 ± 1.2 | 5 |
| Hsapi-Kv2.1 | 1.04 ± 1.2 | 13.5 ± 0.71 | 6 |
| Hsapi-Kv2.1 + Hsapi-Kv8.2 | 10.8 ± 4.2 | 13.8 ± 2.9 | 6 |
| Hsapi-Kv2.1 + Oanat-Kv8.3 | 6.2 ± 3.4 | 12.0 ± 1.1 | 6 |
| Hsapi-Kv2.1 + Hsapi-Kv9.3 | -5.4 ± 6.1 | 16.3 ± 1.6 | 6 |
| Hsapi-Kv2.1 + Lchal-Kv9.4 | -4.7 ± 2.1 | 14.9 ± 1.6 | 6 |
| Drotu-Kv3.1 | 11.4 ± 1.8 | 12.9 ± 0.45 | 6 |
| Drotu-Kv3.1 + Drotu-Kv3.5 | 13.4 ± 1.6 | 8.48 ± 1.4 | 6 |
| **<u>Steady State Inactivation (SSI) analyses</u>** |  |  |  |
| **Channel** | **V<sub>50</sub> (mV)** | **Slope (mV)** | **n** |
| Hsapi-Kv12.1 | -15.9 ± 1.5 | 5.92 ± 0.36 | 7 |
| Hsapi-Kv2.1 + Hsapi-Kv8.2 | -26.9 ± 1.5 | 6.54 ± 0.26 | 7 |
| Hsapi-Kv2.1 + Hsapi-Kv9.3 | -42.2 ± 2.1 | 5.84 ± 0.86 | 6 |
| Hsapi-Kv2.1 + Oanat-Kv8.3 | -22.6 ± 1.9 | 5.39 ± 0.66 | 6 |
| Hsapi-Kv2.1 + Lchal-Kv9.4 | -35.4 ± 2.4 | 9.11 ± 1.1 | 6 |

\*All measurements show mean ± S.D.

The most conserved functional feature of KCNQ channels is a phosphatidylinositol 4,5-bisphosphate (PIP2) requirement for activation (Wang et al., 1998; Zaydman et al., 2013; Li et al., 2015b). We therefore co-expressed Kv7.1L with the voltage-dependent phosphoinositide phosphatase Ciona VSP (Murata et al., 2005) to test its PIP2-dependence. We used human Kv7.2 and Kv2.1 as positive and negative controls, respectively. VSP co-expression inhibits activation of Kv7 currents in oocytes in response to repeated depolarizations (Kruse et al., 2012; Zaydman et al., 2013), but Kv2.1 activation is resistant to VSP-induced PIP2 depletion (Kruse et al., 2012; Delgado-Ramírez et al., 2018). Both Kv7.2 and Kv7.1L (but not Kv2.1) currents were significantly reduced after 10 depolarizes pulses in the presence of Ciona VSP, confirming Kv7.1L retains the classic PIP2-dependence of KCNQ channels (Fig. 3D,E).

### EAG family

A BI phylogeny of the animal EAG family is shown in Fig. 4 with support from both the BI and ML phylogenies overlaid at key nodes. The gene family sequence alignment and fully annotated tree files are provided as Data S13-15. We used ctenophore EAG channels as the outgroup because they diverged prior to the origin of the Eag, Elk and Erg subfamilies in the cnidarian bilaterian ancestor (Martinson et al., 2014; Li et al., 2015c; Jegla et al., 2024; Simonson et al., 2025). Both trees support the presence of a single ancestral gene for each subfamily in the common bilaterian ancestor, the protostome ancestor, the deuterostome ancestor and the vertebrate ancestor. There are duplications of the Erg ancestor in echinoderms and hemichordates in the deuterostome lineage and chelicerates and mollusks in the protostome lineage (Table 1). Chelicerates also have a duplication in the Eag subfamily, and there are additional species-specific duplications in some protostomes for all three subfamilies. Cnidarians also have duplications in the Erg subfamily and the Eag subfamily (hydrozoans only) (Martinson et al., 2014; Li et al., 2015c; Lara et al., 2023).

**Figure 4.**
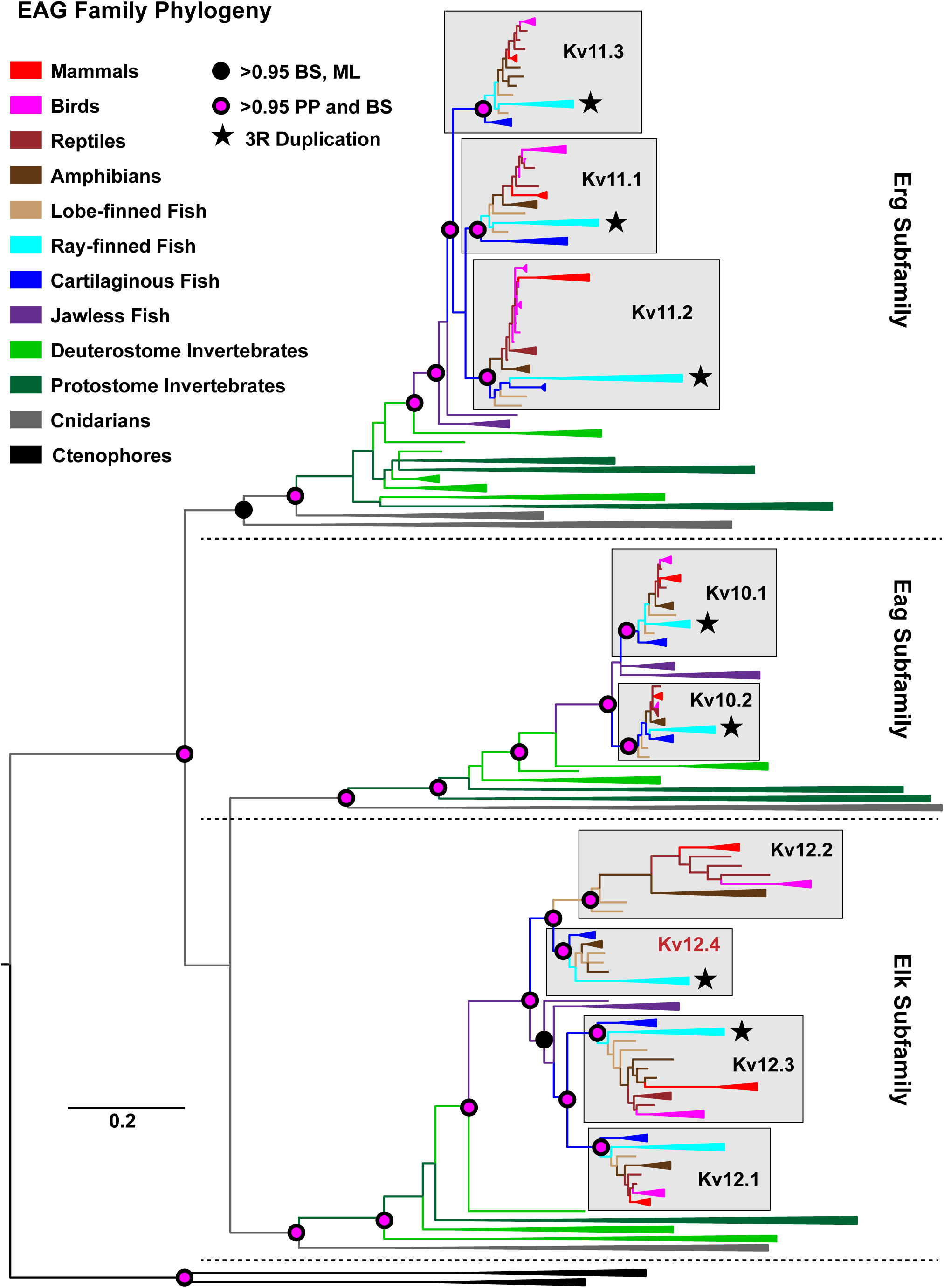
Phylogenetic analysis of the EAG gene family. Bayesian inference phylogeny of EAG family using a 596-site alignment of 447 channels representing a broad diversity of animals. Taxa are colored by phylogenetic group and clades with multiple sequences from the same group are collapsed as in Fig. 2. The tree is rooted for display using ctenophore EAG channels as an outgroup. Three recognized gene subfamilies (Eag, Erg and Elk) are labeled and separated with dotted lines, and ortholog groups containing the 8 human EAG family channels are shaded gray and labeled according to the IUPHAR nomenclature. We named a 9^th^ ortholog group in the Elk subfamily Kv12.4. Stars in ortholog groups indicate candidate 3R duplications in teleost fish and the scale bar for branch length indicates substitutions/site. Statistical support at select nodes (> 0.95) in this phylogeny and a 2^nd^ maximum likelihood (ML) phylogeny based on the same alignment are indicated with colored circles.

Seven of eight human EAG family channels belong to pan-gnathostome clades with strong statistical support in both phylogenies, including both Eag subfamily channels (Kv10.1, Kv10.2), all three Erg subfamily channels (Kv11.1-3) and the Elk subfamily channels Kv12.1 and Kv12.3. Kv10.1 and Kv10.2 duplicated from a single ancestor within gnathostomes after the split from cyclostomes and are therefore likely 2R ohnologs. The Erg subfamily has the same overall topology with duplication after the split from cyclostomes, but with three gnathostome orthologs. This suggests 2R duplication accompanied by an additional independent gnathostome-specific duplication, but it isn’t possible to tell which two channels are the original 2R ohnologs. Kv12.1 and Kv12.3 form a putative 2R ohnolog pair with pan-gnathostome distribution and basal pairing with cyclostome sequences, though the inclusion of cyclostome sequences in the ML phylogeny receives only 0.93 bootstrap support. Kv12.2 is present only in Sarcopterygii (tetrapods + lobe-finned fish) and it surprisingly pairs with a 4^th^ Elk channel which we name Kv12.4 which was present in the gnathostome ancestor but has been lost in reptiles, birds and mammals. Only lobe-finned fish and amphibians have both Kv12.2 and Kv12.4. The most parsimonious interpretation of the topology is that Kv12.2 was duplicated from Kv12.4 within Sarcopterygii. Kv12.4 itself is most likely the product of an individual duplication in the stem gnathostome because the Kv12.2/Kv12.4 clade is not paired with cyclostome sequences. We found no evidence for contribution of the 1R genome duplication to the human EAG family because the phylogenies support a single gene for each subfamily at the time of the cyclostome/gnathostome divergence.

Several of the bird species we examined for phylogenetic analysis did not have a Kv12.2 ortholog, but we did find Kv12.2 orthologs in *Apteryx mantelli* (brown kiwi), *Harpia harpyja* (Harpy eagle) and *Grus americana* (whooping crane). Bird genomics is highly advanced and there are now gene predictions from hundreds of high-quality genomes spanning their full phylogenetic diversity. We therefore searched broadly across bird orders to gain insights into the pattern of Kv12.2 loss. We could confidently identify bird Kv12.2 orthologs by homology in BLASTP searches using the three aforementioned bird Kv12.2s as queries because bird Kv12.2 orthologs share > 75% amino acid identity with each other, but < 55% amino acid identity with Kv12.1 and Kv12.3 paralogs. We found Kv12.2 orthologs in only a small subset of bird species within the following clades: Paleognathae (flightless ratites including kiwis), Gruiformes (cranes, rails and allies), Accipitriformes (hawks and eagles) and Aequornithes (loons, penguins, storks, pelicans and allies) (Fig. 5). In addition, the clade Strisores, which includes hummingbirds swifts and nightjars, has Kv12.2 in only one species: the South American oilbird, which has been assigned to its own order (Stiller et al., 2024). Explaining the Kv12.2 distribution we found remarkably requires 10 independent losses over the 100+ million years of bird evolution (Fig. 5).

**Figure 5.**
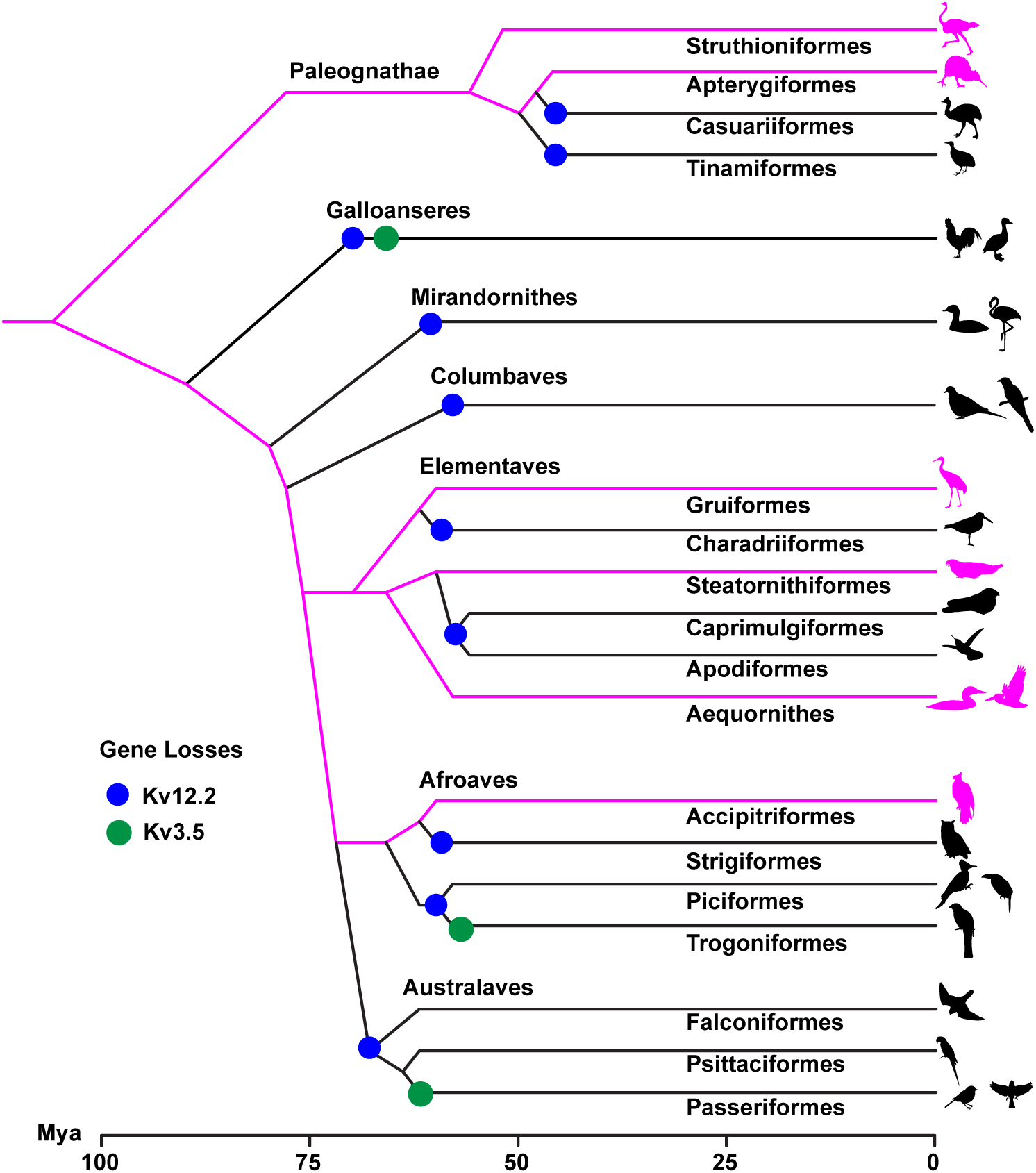
Loss of Kv12.2 and Kv3.5 in birds. Schematic phylogeny of major bird orders based on the phylogenetic relationships determined from comparison of hundreds of bird genomes (Stiller et al., 2024) with approximate divergence times in millions of years ago (Mya) taken from TimeTree 5 (Kumar et al., 2022). Bird orders are labeled near branch tips while higher order classifications are labeled near branch bases as needed. Lineages with Kv12.2 are highlighted in magenta, and 10 independent losses of Kv12.2 necessary to explain the distribution pattern are indicated with blue circles. Green circles mark three losses of the Shaker family channel Kv3.5 but the Kv3.5 distribution pattern is not highlighted on branches. Bird icons used to label branches are open-source images from the PhyloPic.org repository.

Teleost fish retain 3R duplications for 7 of the 8 gnathostome EAG family channels, with Kv12.1 the only exception. The teleost ancestor likely had 15 EAG family channels and the extant species we analyzed here have 13-15 of these channels, though only the *Ictalurus punctatus* has the full teleost set. Zebrafish, a model organism with a highly curated genome, has 13 EAG family genes with presumptive 3R duplications for Kv10.1, Kv10.2, Kv11.1, Kv11.2 and Kv12.3. 3R duplications of Kv11.3 and Kv12.4 appear to have been lost in zebrafish. We have lower confidence in differentiating losses from missing data in the other three teleost species because their genomes are likely to be less complete. ZElk, a zebrafish Kv12 channel which has previously been used for structural and biophysical analyses (Haitin et al., 2013; Dai and Zagotta, 2017; Dai et al., 2018), turns out to be zebrafish Kv12.4 and thus has no ortholog in mouse and human.

We expressed the axolotl Kv12.4 ortholog (Amexi-Kv12.4) and compared it to human Kv12.1-12.3 currents recorded under the same conditions. Example current traces for Kv12.1-12.4 recorded from Xenopus oocytes are shown in Fig. 6A and GV curves are shown in Fig. 6B with Boltzmann fit parameters given in Table 2. Kv12.1 and Kv12.3 show little inactivation, but Kv12.2 and Kv12.4 exhibit varying degrees of fast inactivation at depolarized voltages. The hallmark of human Kv12.1-12.3 channels is a hyperpolarized activation range, though there is a broad spread in V_50_ between the channels, ranging here at pH 7.2 from −83.2 ± 1.5 mV in Kv12.3 to −20.4 ± 3.0 mV in Kv12.2. While Kv12.2 has a more depolarized V_50_ than the others, this results mostly from a low slope (Table 2) and thus activation begins in the same hyperpolarized voltage range as Kv12.1 and Kv12.3. Both Kv12.1 and Kv12.2 have been shown to regulate subthreshold excitability in mammalian neurons and a Kv12.2 mutation has been associated with human epilepsy (Zhang et al., 2010; Hermanstyne et al., 2023; Bauer et al., 2025a; Bauer et al., 2025b). The V_50_ of Amexi-Kv12.4 was not significantly different human Kv12.2 (*p* = 0.113, t-test, n = 5-6), but the GV does have a significantly steeper slope (*p* < 0.001, t-test, n=5-6) leading to reduced Kv12.4 activation at hyperpolarized potentials. However, Kv12.4 current is consistently present at voltages as low as −60 to −70 mV, so it could still influence subthreshold excitability. Amexi-Kv12.4 has significantly faster and more steeply voltage-dependent deactivation than the other vertebrate Elk channels (Fig. 6C) (Anova with Benjamini-Hochberg corrected post-hoc pairwise testing, p < 0.01, n = 5-8), but we did not examine whether this is responsible for the GV shift. We confirmed the presence of inactivation in Amexi-Kv12.4 and characterized its properties with a reactivation protocol (Fig. 6D). Inactivation was limited to depolarized voltages and was significantly slower and less complete than inactivation in Hsapi Kv12.2. However, inactivation with similar properties is conserved in the zebrafish Kv12.4 ortholog zElk along with the comparatively depolarized GV and rapid deactivation (Dai and Zagotta, 2017), so it may nonetheless have functional significance. Kv12 channel activation kinetics and V_50_ are highly sensitive extracellular pH, Zn^2+^ and Cd^2+^ via interaction with conserved acidic residues in the extracellular aqueous pocket of the VSD (Zhang et al., 2009; Kazmierczak et al., 2013), but our recordings and previous zElk recordings were made at the same pH and therefore shared properties likely represent true functional conservation. These results suggest that the loss of Kv12.4 in reptiles, birds and mammals may represent the loss of a unique Kv12 conductance.

**Figure 6.**
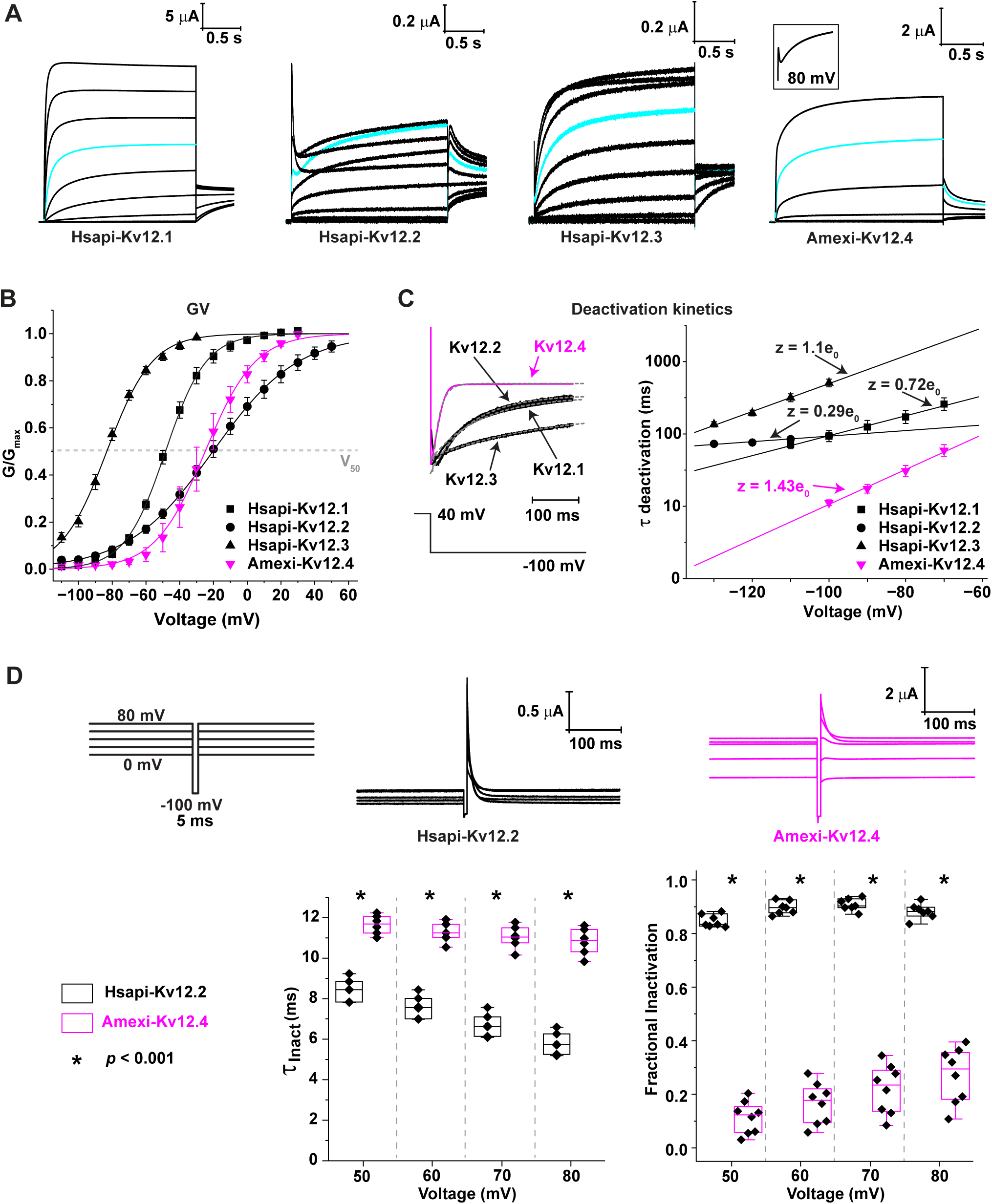
Functional comparison of vertebrate Kv12.1-4 paralogs. (A) Families of currents recorded from oocytes expressing Hsapi-Kv12.1-3 and Amexi-Kv12.4. Kv12.1-3 currents were recorded in response to depolarizing voltage sweeps ranging from −100 mV to 60 mV in 20 mV increments with the 0 mV sweep highlighted in cyan. Tails were recorded at −40 mV. Kv12.4 currents were not initially recorded at high voltages as this was unnecessary for GV analysis, so 20 mV is top sweep displayed. We later independently recorded currents at high voltage to look for signs of inactivation and the inset shows an example Amexi-Kv12.4 trace recorded at 80 mV with inactivation displayed according to the same scale bar. (B) Normalized GV curves for Kv12.1-Kv12.4 measured from isochronal tail currents with data shown as mean ± S.D. Curves show single Boltzmann fits using average V_50_ and slope parameters determined from fits of individual oocytes (Table 2). (C, right) Normalized tail current example traces recorded at −100 mV (after a 1 s test pulse to 40 mV) overlaid with single exponential fits (dotted lines) that were used to determine deactivation time constants. (C, left) Semi-log plots of deactivation time constant vs. voltage fit with single exponentials to estimate the elemental charge associated deactivation (z). (D) The inactivation properties of Kv12.2 and Kv12.4 were determined using a protocol in which two depolarizing pulses were separated by a 5 ms repolarization to allow recovery from inactivation (rapid in Kv12s) without significant deactivation. Fractional inactivation and inactivation time constant can then be measured in the 2^nd^ pulse (Trudeau et al., 1999; Zhang et al., 2010). Example current traces for Kv12.2 and Kv12.4 recorded with this protocol are shown in the top row and clipped to magnify the region surrounding the recovery step. Plots of inactivation time constant and fractional inactivation vs. are shown at the bottom. Boxes display middle quartiles and median, whiskers mark the observed data range and data from individual oocytes are shown with black diamonds. Time constants and fractional inactivation differed significantly between Kv12.2 and Kv12.4 at all voltages (p < 0.0001, t-test).

### Shaker Kv1 subfamily

We split the Shaker family into subfamilies for phylogenetic analysis for computational efficiency because of the large number these channels in the animal species we examined. Shaker family channel sequences are shorter than KCNQs and EAGS, so the split also helped us increase the length of alignments for some of the subfamilies. A BI phylogeny of the Kv1 subfamily is shown in Fig. 7; see Data S16-S18 for the alignment file and BI and ML trees. All 8 human Kv1s fall into pan-gnathostome ortholog clades. 6/8 clades receive consensus support in both phylogenies, but the Kv1.3 clade only received > 0.95 support in BI, while the Kv1.6 was supported at > 0.95 only in ML. Because the full Kv1 alignment was only 330 positions, we made a vertebrate-only alignment (excluding a single divergent cyclostome clade that branched with invertebrate sequences), but this only increased the alignment length to 389 positions. We nevertheless re-ran the phylogenies with BI and ML and the alignment and trees are provided as Data S19-S21. The only substantive change in these vertebrate-only phylogenies was to establish consensus support for the pan-gnathostome Kv1.3 and Kv1.6 ortholog clades shown in Fig. 7.

**Figure 7.**
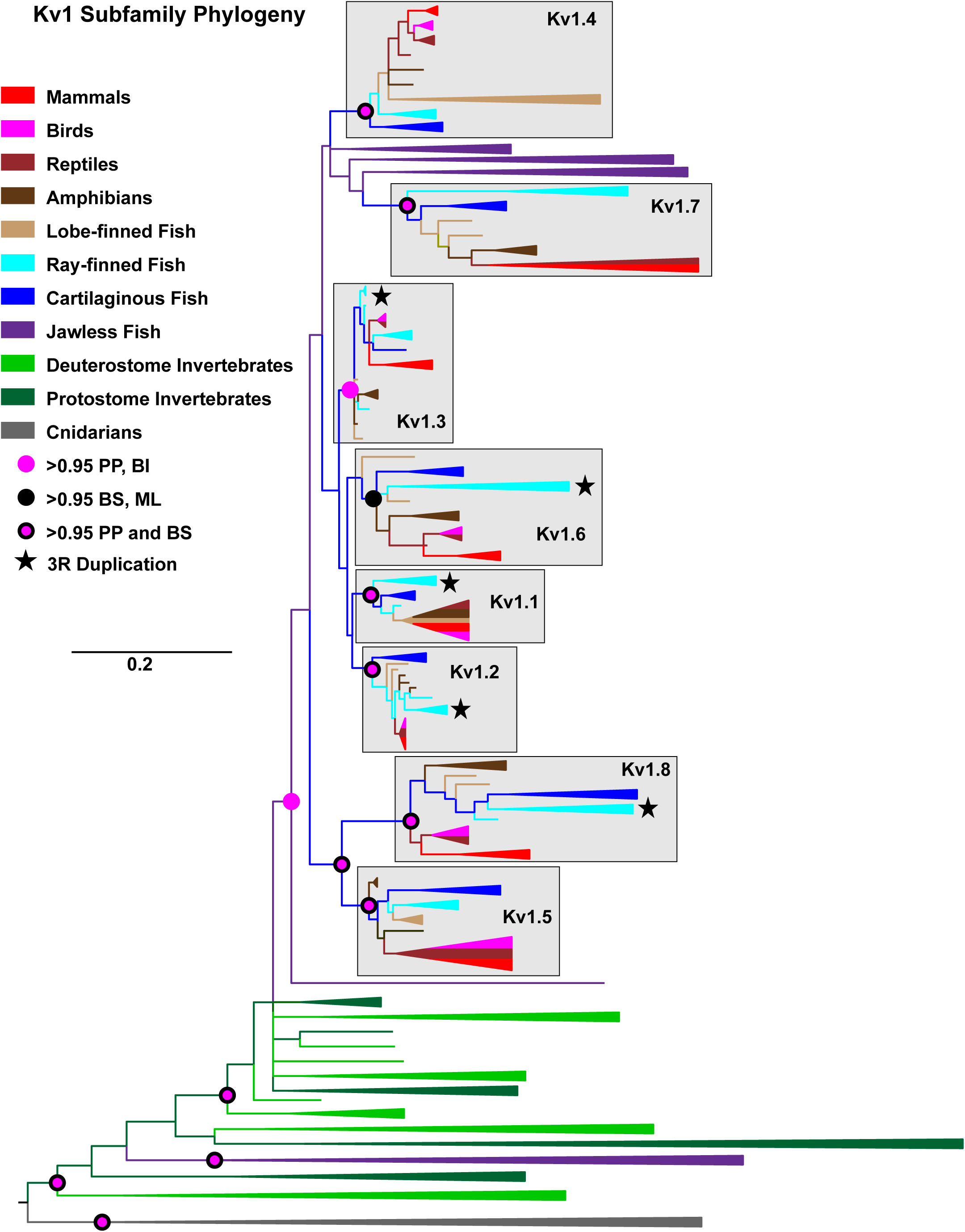
Phylogenetic analysis of the Kv1 subfamily. Bayesian inference phylogeny of the Kv1 subfamily of the Shaker gene family based on a 330-site alignment of 356 channels spanning the animal phylogeny. Taxa and clades are displayed as in Figures 2 and 4 and colored according to the legend. Split colors are used for collapsed clades containing intermixed sequences from multiple groups; the extra clade width of mixed clades was used only to show the multiple colors more clearly and is not scaled to the number of taxa. The tree is rooted with cnidarians as an outgroup for display. The 8 shaded ortholog groups all contain a human sequence and use IUPHAR names. Stars indicate putative 3R duplications, node support for key clades in this phylogeny and a ML phylogeny using the same alignment is marked according to the legend, and the scale bar indicates substitutions/site.

The relationship to cyclostome channels was not further resolved and the strict interpretation of the phylogenies is that the vertebrate ancestor had a single Kv1 gene that was independently duplicated in cyclostomes and gnathostomes, meaning that the 1R genome duplication did not contribute to our Kv1 channel set. Our 8 Kv1 channels map to just 4 chromosomal loci as there are two arrays of three tandem Kv1 subfamily genes in mammals (Albrecht et al., 1995; Street and Tempel, 1997): Kv1.3-Kv1.2-Kv1.8 (human chromosome 1) and Kv1.6-Kv1.1-Kv1.5 (human chromosome 12). Cyclostomes do not have these tandem Kv1 arrays but all gnathostomes do. The logical explanation is that the two arrays were duplicated in 2R from an ancestral 3 gene array created by local duplications in the gnathostome common ancestor. The six Kv1 channels in the two arrays are therefore likely 2R ohnologs. Whether Kv1.4 and Kv1.7 are also 2R ohnologs or derived from independent duplications in the gnathostome lineage is unclear because they do not group together with statistical support. Our analysis predicts the teleost ancestor had 13 Kv1 genes including 4 Kv1 arrays, though one array missing a Kv1.5 co-ortholog. Zebrafish retains the full ancestral teleost Kv1 subfamily set.

Kv1 channels have also been extensively duplicated in bilaterian invertebrates; we find independent duplications in tunicates, cephalochordates, lophotrochozoans (mollusks, annelids and platyhelminthes) and chelicerates (Table 1). Only insects, nematodes and echinoderms lack Kv1 subfamily duplications. The largest Kv1 subfamilies are found in cnidarians and cephalochordates rather than vertebrates. 12 ancestral Kv1 lineages are widely conserved in cnidarians (Lara et al., 2023) and the model species *Nematostella vectensis* (starlet sea anemone) has 20 Kv1 channels including as many as 14 silent subunits (Jegla et al., 2012).

### Shaker Kv2 Subfamily

We performed BI and ML phylogenetic analysis on the entire Kv2 subfamily including the vertebrate silent subunits (Kv5,6,8,9) and the BI tree is shown in Fig. 8A (alignment and tree files are Data S22-24). There is consensus phylogenetic support for pan-gnathostome clades of all 10 human silent subunits and for the existence of two additional silent subunits, named Kv8.3 and Kv9.4 here, that have been lost in humans and mice. Kv8.3 is broadly present in gnathostomes but was lost in mammals after the divergence of the egg-laying monotremes.

**Figure 8.**
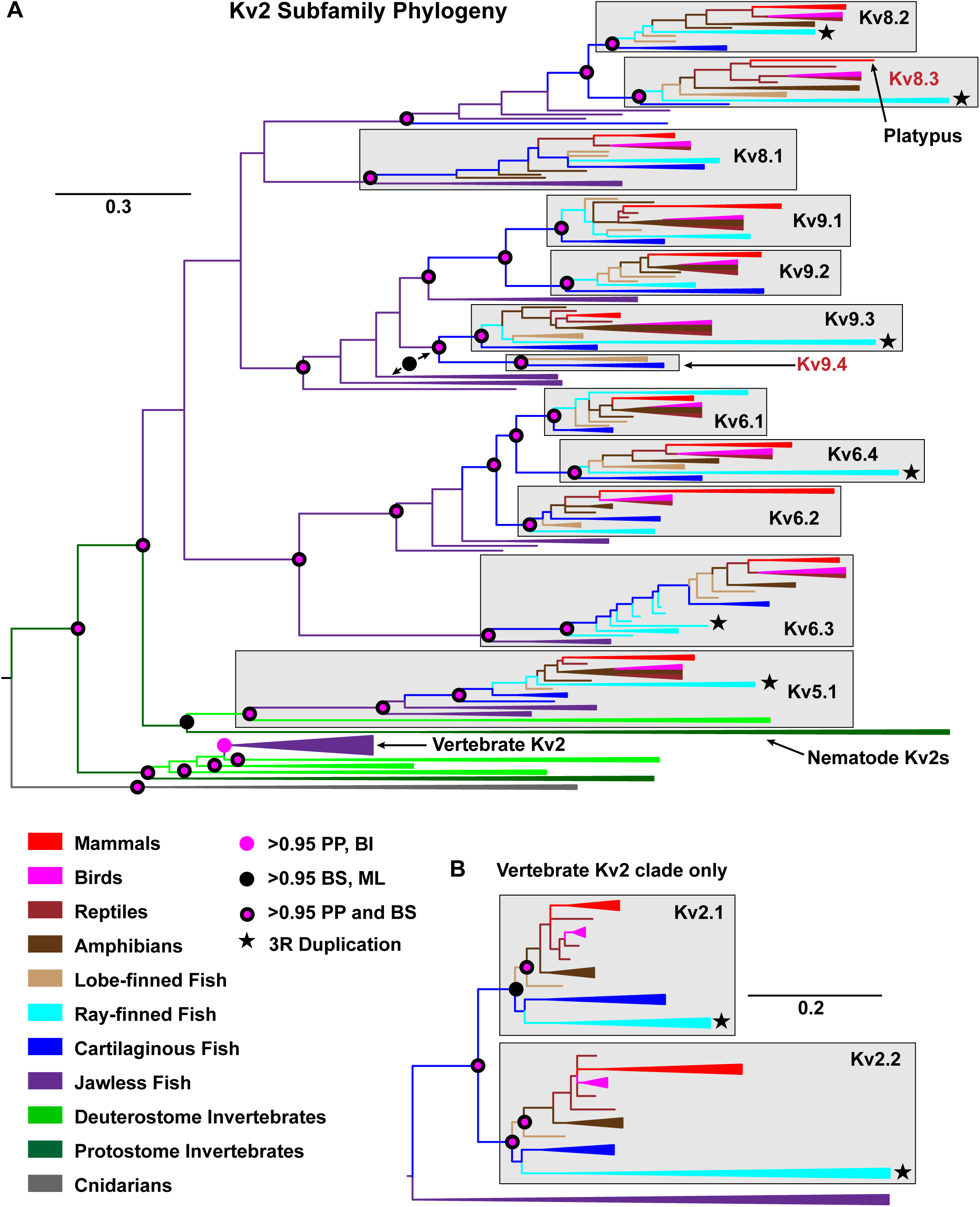
Phylogenetic analysis of the Kv2 subfamily. (A) Bayesian inference phylogeny of the Shaker gene family’s Kv2 subfamily made using a 393-site alignment of 509 sequences and rooted between cnidarian and bilaterian clades for display. Taxa are colored according to the legend, and the width of collapsed clades is not proportional to sequence number. Collapsed clades with multiple colors include intermixed sequences from multiple groups. 12 vertebrate silent subunit ortholog groups are shaded and labeled according to the IUPHAR nomenclature extended to cover the new channels Kv8.3 and Kv9.4 which are absent in humans and mice. Statistical support at selected nodes important for interpretation is indicated according to the legend and includes support from a parallel ML phylogeny that was run on the same alignment. Stars mark 3R duplications in teleosts and the scale bar is in substitutions/site. (B) Bayesian inference phylogeny of 74 vertebrate Kv2 α-subunits based on a 681-site alignment. Pan-gnathostome Kv2.1 and Kv2.2 clades (shaded) can be resolved with this extended alignment, though only ML supports inclusion of cartilaginous fish and ray-finned fish with > 0.95 support in the Kv2.1 clade. BI support for a pan-gnathostome Kv2.1 clade was 0.89 PP.

Kv9.4 is present only in cartilaginous fish and lobe-finned fish implying independent losses in tetrapods and ray-finned fish. There are 6 gnathostome/cyclostome silent subunit clades with consensus support: Kv8.2/Kv8.3, Kv8.1, Kv9.1/Kv9.2, Kv6.1/Kv6.2/Kv6.4, Kv6.3 and Kv5.1. In addition, ML supports a 7^th^ cyclostome/gnathostome Kv9.3/Kv9.4 clade. This pattern indicates extensive silent subunit expansion in the vertebrate common ancestor prior to 1R, followed by additional 1R duplications in the Kv6 and Kv9 clades. The phylogenies don’t statistically support a unified Kv8 clade, so we can’t assume the Kv8.1 and Kv8.2/8.3 clades also derive from a 1R duplication; both clades might have already been present in the vertebrate common ancestor. The phylogenies support four 2R duplications to further expand the Kv2 subfamily silent subunits in gnathostomes, and one additional independent duplication in the stem gnathostome lineage to add a third gene to the Kv6.1/Kv6/2/Kv6/4 clade. 2R duplications were not retained for the Kv6.3, Kv8.1, and Kv5.1 clades. Our phylogenetic analyses confirm past results that Kv5.1 is the oldest Kv2 subfamily silent subunit with an origin in the chordate ancestor of tunicates and vertebrates (Pisupati et al., 2018; Jegla and Simonson, 2023). All vertebrate silent subunits fall into a single clade with consensus support in BI and ML, so Kv6, Kv8 and Kv9 channels most likely arose via duplication of an ancestral Kv5-like gene.

Inclusion of the silent subunits in the Kv2 subfamily trees reduced alignment length to the point that there weren’t enough informative sites to resolve the highly conserved vertebrate Kv2.1 and Kv2.2 clades. We therefore ran a 2^nd^ set of phylogenetic analyses on just vertebrate Kv2 α-subunits which allowed us to increase alignment length from 393 to 681 positions (Fig. 8B, Data S25-S27). This 2^nd^ analysis provides consensus support for a pan-gnathostome Kv2.2 clade and ML also supported a pan-gnathostome Kv2.1 clade. The BI phylogeny also had a pan-gnathostome Kv2.1clade, but with only 0.89 PP for inclusion of cartilaginous and ray-finned fish sequences. Alternative explanations for the BI tree topology would require invocation of multiple independent duplication/loss events. Kv2 α-subunits were extensively duplicated in cyclostomes, but there is consensus support for split of Kv2.1 and Kv2.2 after divergence from cyclostomes, making them a likely 2R ohnolog pair.

There are apparent 3R duplications for Kv2.1, Kv2.2, Kv5.1, Kv6.3, Kv6.4, Kv8.2, Kv8.3 and Kv9.4 in teleost fish based on the presence of co-orthologs in multiple teleost species. Total Kv2 subfamily gene counts in the teleost species we examined ranging from 17 in zebrafish to 22 in european eel (*Anguilla anguilla*). Within protostomes, we found Kv2 duplications only in chelicerates and nematodes (Wei et al., 1996), and the Kv2 subfamily is represented by a single gene in cnidarians (Li et al., 2015b; Lara et al., 2023). Nematode Kv2s are highly divergent and gravitate into the vertebrate silent subunit clade next to Kv5s in our phylogenies. We think this most likely represents a case of long-branch attraction (Bergsten, 2005) since a true evolutionary link between nematode Kv2s and vertebrate Kv2 subfamily silent subunits would require invoking a large number of independent gene loss events across the bilaterian phylogeny.

Most invertebrate Kv2 subfamily channels have long C-terminal cytoplasmic domains, so we looked for conservation with the C-termini of vertebrate Kv2 orthologs which contain a non-canonical FFAT domain (ncFFAT, also know as the PRC domain). This domain mediates direct interaction between Kv2.1 and Kv2.2 and ER-resident VAP proteins to give mammalian Kv2 channels a non-conducting role in organizing plasma membrane ER contacts (Fox et al., 2015; Johnson et al., 2018; Kirmiz et al., 2018a; Kirmiz et al., 2018b). We found evidence for widespread conservation of the ncFFAT/PRC domain in Kv2 channels across Bilateria (Fig. 9A). This interaction domain was previously proposed to be vertebrate specific primarily because it is absent in the original published short isoform of Drosophila Shab. The site is however present in long splice variants of Drosophila Shab and many other invertebrate Kv2 sequences. Fig. 9B shows the distribution of Kv2 channels with a putative ncFFAT/PRC in animals overlaid on the BI phylogeny from Fig. 8A. The most parsimonious interpretation of the distribution is that the ncFFAT/PRC arose in the ancestral bilaterian Kv2 channel but has been lost as many as eight different times in bilaterians, including in the vertebrate Kv2 subfamily silent subunits. Some protostomes, including some arthropods, have versions of Kv2 channels with and without the ncFFAT/PRC. In Drosophila, this is done by alternative splicing, but in chelicerates it is done with different genes. We propose the non-conducting role of Kv2 channels in organization of plasma membrane-ER contacts most likely evolved in the bilaterian common ancestor, much earlier than previously thought. Similarly, giant ankyrin isoforms that organize the axon initial segment were originally thought to be vertebrate specific but broader phylogenetic analyses identified giant ankyrins as pan-bilaterian and showed a conserved role for them in establishing an axon initial segment diffusion barrier in *Drosophila* (Jegla et al., 2016).

**Figure 9.**
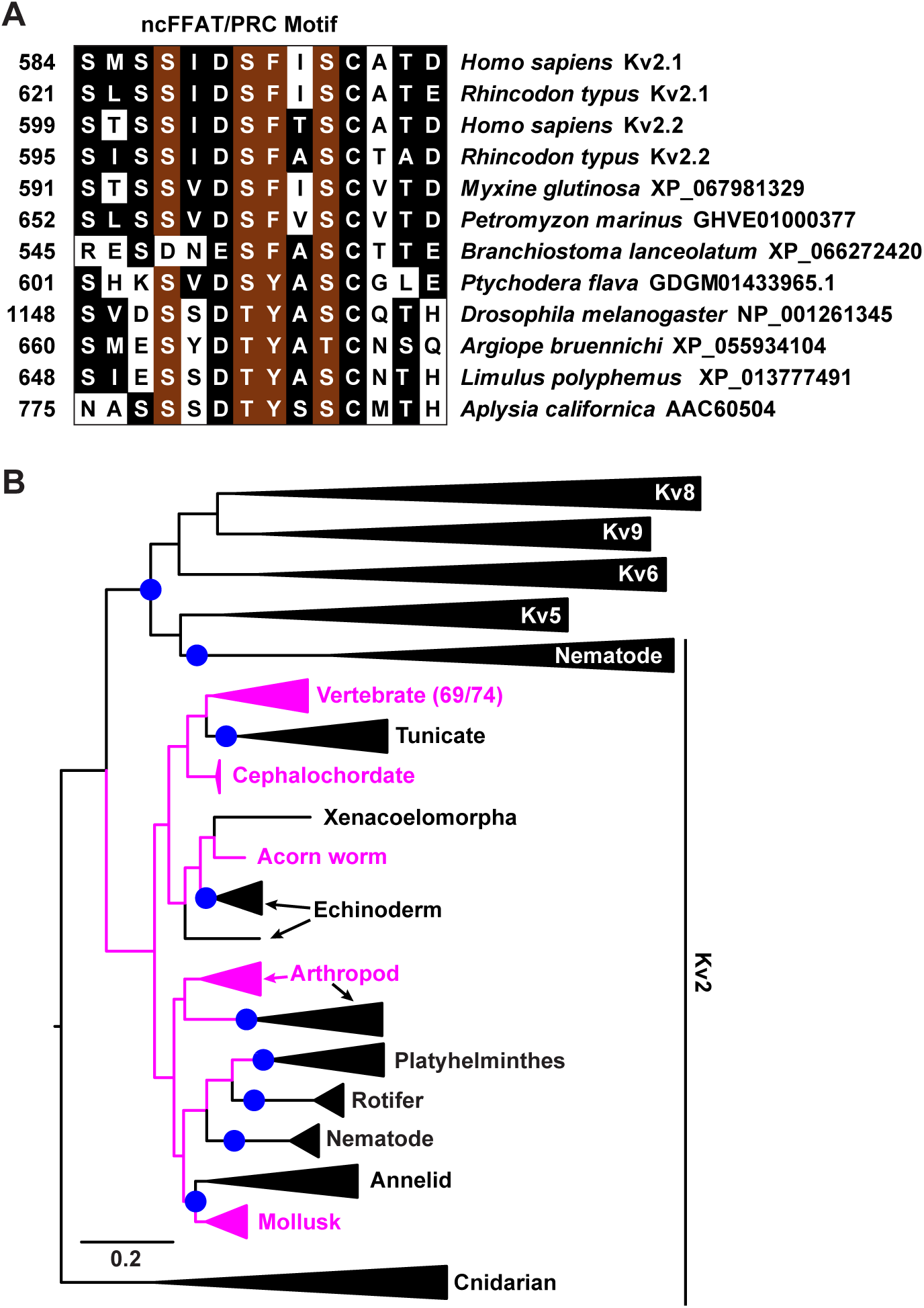
Evolutionary conservation of a VAP binding site in bilaterian Kv2 α-subunits. (A) Sequence alignment of the ncFFAT/PRC motif that mediates Kv2-VAP interactions associated with the formation of plasma membrane-ER contacts in mouse neurons. Sequence positions are given on the left and species labels on the right. Positions conserved identically or with conservative substitution in more than half the sequences are shaded and red shading marks positions known to have a role in Kv2-VAP association. (B) Overlay of ncFFAT/PRC presence (magenta) on the BI phylogeny from Fig. 8A. The ncFFAT/PRC motif can be traced to the ancestral bilaterian Kv2 channel but has been lost (blue dots) at least 9 independent times during Kv2 subfamily evolution. 69/74 vertebrate Kv2 α-subunits had the ncFFAT/PRC motif, but we did not score missing motifs within this clade as true losses because we did examine the possibility of splice variation in gene predictions.

We expressed Kv8.3 and Kv9.4 orthologs from platypus (*Ornithorhynchus anatinus*, Oanat-Kv8.3) and Coelacanth (*Latimeria chalumnae*, Lchal-Kv9.4) in Xenopus oocytes to examine their functional properties. Platypus and Coelacanth represent the closest living relatives to humans that still have these genes. Since silent subunits require assembly with Kv2.1 or Kv2.2, we co-expressed Oanat-Kv8.3 and Lchal-Kv9.4 with human Kv2.1 (Hsapi-Kv2.1) to assess their function. We also compared the heteromeric currents to Hsapi-Kv2.1/Hsapi-Kv8.2 and Hsapi-Kv2.1/Hsapi-9.3 heteromers to assess the evolutionary conservation of functional phenotypes.

Fig 10A shows example traces for Oanat-Kv8.3, Lchal-Kv9.4, Hsapi-Kv2.1 and the four heteromeric subunit combinations. Current size comparisons are shown in Fig. 10B. Oanat-Kv8.3 and Lchal-Kv9.4 are indeed silent subunits because they do not functionally express as homomultimers but can co-assemble with Kv2.1 based on 1) significant reductions in total current size we observed upon co-expression (t-test, *p* < 0.05) and 2) significant differences between the biophysical properties of homomeric Kv2.1 currents and K^+^ currents in oocytes expressing Kv2.1 along with these silent subunits. Reductions in current size for silent/α-subunit co-expression compared to α-subunit controls are expected if stoichiometry restrictions reduce overall assembly efficiency (Pisupati et al., 2018). Note however that the degree of suppression varies with subunit ratios, so differences between heteromeric currents should only be made with a broad titration of expression ratios which we did not do here. Therefore, we can’t conclude that the difference in heteromeric current size we see here are biological – they could just represent differences in α/s subunit ratios. The GV V_50_ shifts induced by Kv8s and Kv9s were small and not significantly different between Oanat Kv8.3 and Hsapi Kv8.2 or between Lchal Kv9.4 and Hsapi Kv9.3 (Fig. 10C). However, the Kv2.1-Kv8.x and Kv2.1-Kv9.x V_50_s were significantly different because the shifts were in opposite directions: positive for the Kv8s and negative for the Kv9s. Both Kv8s and Kv9s shifted the steady state inactivation currents in a hyperpolarizing direction, but the size of the shift was significantly bigger for the Kv9s (Fig. 10D). All four silent subunits also significantly slowed both deactivation and activation (Fig. 10E,F), but the slowing of deactivation for Kv9s was significantly larger compared to Kv8s. Statistical comparisons of V_50_s and kinetics were made with one-way ANOVA analyses followed by Benjamini–Hochberg corrected post hoc pairwise testing (n.s., *p* > 0.05; significant, *p* < 0.01 and n = 6-7 for V_50_s, *p* < 0.05 and n = 6-7 for kinetics). There were also some significant differences in the biophysical properties for Hsapi-Kv8.2 vs. Oanat-Kv8.3 and for Hsapi-Kv9.3 vs. Lchal-Kv9.4, but these were not large as the differences between Kv8s and Kv9s. We think it is premature to assign any functional significance to differences that stem from smaller effects for Oanat-Kv8.3 or Lchal-Kv9.4 because these could simply stem from less-efficient cross-species assembly. Whole oocyte currents will include contributions from both the α/s-subunit heteromers and α-subunit homomers and one minimally needs an ∼3-fold excess of silent subunit to obtain majority heteromeric currents (Pisupati et al., 2018). Assembly efficiency could therefore influence heteromer percentage in addition to current size, and we did indeed need to use higher silent/α-subunit expression ratios to observe heteromeric currents for Oanat-Kv8.3 and Lchal-Kv9.4 (50:1) compared to Hsapi-Kv8.2 and Hsapi-Kv9.3 (3-5:1).

**Figure 10.**
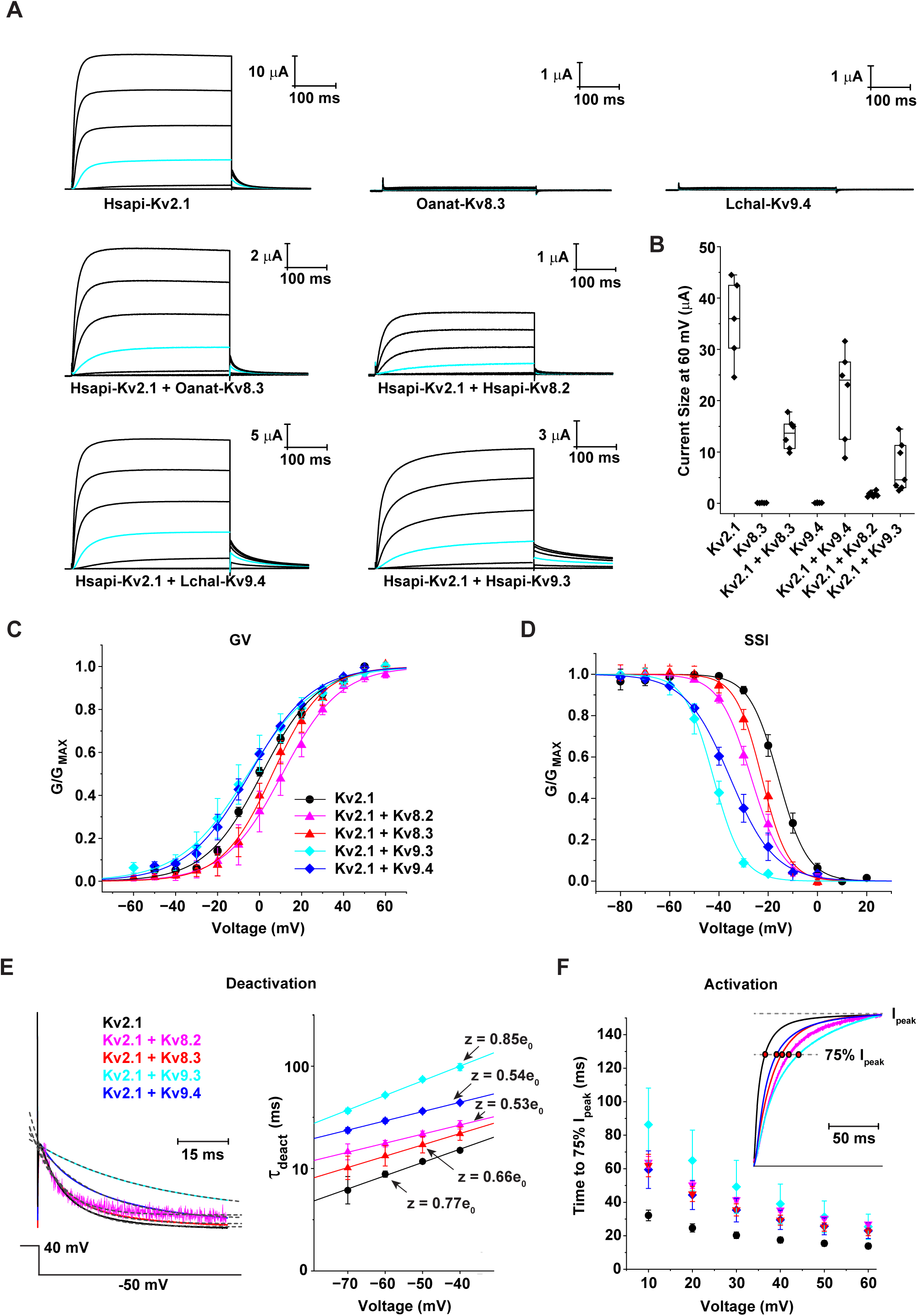
Functional characterization of Kv8.3 and Kv9.4. (A) Example current traces recorded from oocytes expressing the indicated Kv2/Kv8/Kv9 orthologs. Currents were elicited with depolarizing voltage steps ranging from −60 mV to 60 mV in 20 mV increments with tail currents recorded at −40 mV; the 0 mV current traces are highlighted in cyan to facilitate comparison and scale bars are given for current size and time. (B) Peak current size at 60 mV for the specified channel combinations. Boxes show median and middle quartiles, whiskers mark the data range and data points (black diamonds) show measurements from individual oocytes. (C,D) Normalized GV and steady state inactivation (SSI) curves for Hsapi-Kv2.1 expressed alone and in combination with Hsapi-Kv8.2, Oanat-Kv8.3, Hsapi-Kv9.3 and Lchal-Kv9.4. Data points were normalized for display and show mean ± S.D. Curves are single Boltzmann fits using the average V_50_ and slope values determined from individual oocytes (Table 2). (E) Comparison of deactivation time constants for Kv2.1 and Kv2.1-silent subunit combinations. The left panel shows normalized examples of tail currents recorded at −50 mV after a +40 mV test step overlaid with single exponential fits (dotted gray curves). The right panel shows a semi-log plot of deactivation time constants vs. voltage with single exponential fits used to determine the charge associated with deactivation (z). (F) Plot of 75% activation time vs. voltage for Kv2.1 alone and in combination with Kv8 and Kv9 silent subunits. The inset shows an overlay of normalized example current traces for each condition recorded at 40 mV to illustrate the measurement method. Data points in E and F show mean ± S.D. of n = 6 measurements.

### Shaker Kv3 Subfamily

The ML phylogeny of the animal Kv3 subfamily is shown in Fig. 11A, and the alignment plus BI and ML phylogeny files are Data S28-S30. The Kv3.1-3 clades are clearly pan-gnathostome with strong statistical support with both BI and ML, though Kv3.3 has been lost in birds. The Kv3.4 clade has sequences from all gnathostome clades but does not receive basal statistical support. Most interestingly we found evidence for a highly divergent 5^th^ Kv3 channel we named Kv3.5 in ray-finned fish and most tetrapods. Kv3.5 branches from within the Kv3.4 clade in ML, suggesting it could have arisen from a Kv3.4 duplication in the ancestor of bony vertebrates (Osteichthyes) after the split from cartilaginous fish. Kv3.5 has a basal position within the vertebrate Kv3 clade in the BI phylogeny (Fig. 11B), so we re-ran the phylogenetic analysis using only vertebrate Kv3s to increase alignment length with the hope of gaining better clarity on the Kv3.5 origin (alignment and trees provided as Data S31-33). With the longer alignment, both BI and ML support branching of Kv3.5 from within the Kv3.4 clade (Fig. 11B), which is why we chose to show the original ML alignment in Fig. 11A. Analyses using the longer vertebrate-only alignment also strengthen support for a pan-gnathostome Kv3.4 clade, albeit at only 0.93 PP in BI.

**Figure 11.**
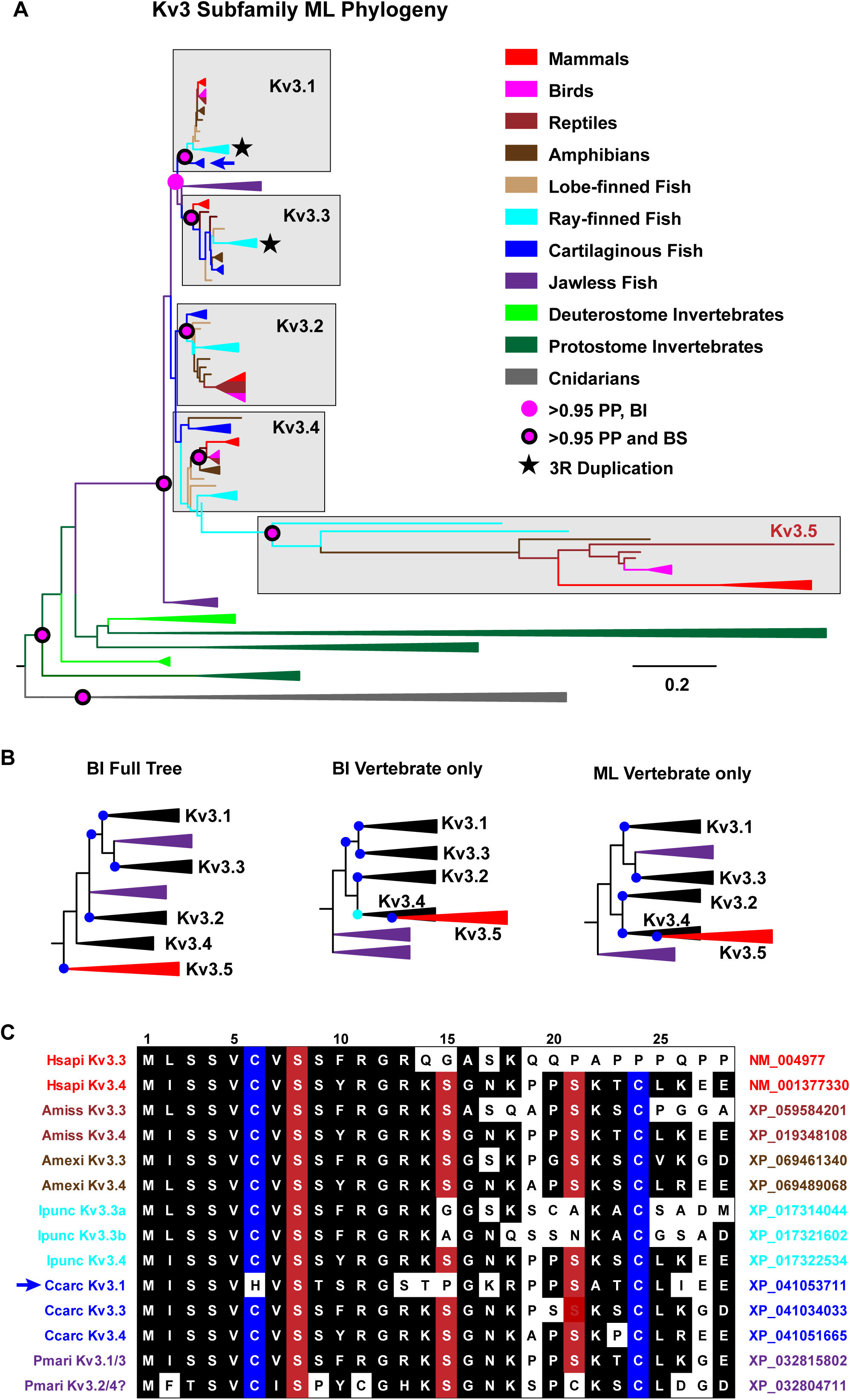
Phylogenetic analysis of the Kv3 subfamily. (A) Maximum likelihood phylogeny of the Kv3 subfamily of the Shaker gene family based on a 235 taxa/373-site alignment and rooted for display between cnidarians and bilaterians. Taxa are colored according to the legend and collapsed clade width is not proportional to sequence number. The scale bar is in units of substitutions/site. Ortholog groups containing human Kv3.1-4 are shaded and labeled, and a new 5^th^ ortholog group we name Kv3.5 branches from within the Kv3.4 clade. Stars in clades indicate likely 3R duplications in teleost fish. Statistical support at key nodes is indicated according to the legend for this phylogeny and a parallel BI phylogeny using the same alignment (B) Schematic diagrams of clade arrangement for Kv3.1-3.5 and two cyclostome Kv3 clades in the aforementioned BI phylogeny (left) and additional BI (middle) and ML (right) phylogenies based on a 158 taxa/484 position vertebrate-only alignment. Clade support of > 0.95 is indicated with blue dots and 0.93 with a cyan dot. Kv3.5 branches from within the Kv3.4 clade in 3 of the 4 phylogenies shown in A and B, and there is support for a pan-gnathostome Kv3.4 clade in the vertebrate-only phylogenies. (C) Amino acid sequence alignment of N-terminal inactivation ball sequences from various vertebrate Kv3 subfamily orthologs. Residues conserved (identical or conservative substitution) in > 50% of the sequences are shaded; blue and red shading is used for cysteines and serine residues implicated in regulation. The arrow highlights a Kv3.1 ortholog from cartilaginous fish with an inactivation ball sequence. Species and ortholog identity are labeled at the left margin and accession numbers are given at the right margin; labels are color coded by phylogenetic group according to the legend in (A). A ruler is provided at the top of the alignment for amino acid position.

We did not find Kv3.5 in lobe-finned fish and it shows a complex pattern of losses in mammals (Fig. 12) and birds (Fig. 5), as determined by comprehensive BLAST searches following the model we used for Kv12.2 in birds. Marsupials, bats, rabbits (Lagomorpha) and rodents are the major Kv3.5+ groups in mammals, and it is conspicuously absent in primates. While Kv3.5 is present in rodents, it has been lost in the rodent clades Histricomorpha, which includes porcupines and guinea pigs, and Myomorpha, which includes rats and mice. Kv3.5 is more widely distributed in birds, but it has been lost in Galloanseraes (chickens, ducks and geese), Trogoniformes (trogons and quetzals) and Passeriformes (perching birds). We did not examine Kv3.5 losses in clades of fish and amphibians, though it wasn’t present in all species we analyzed here.

**Figure 12.**
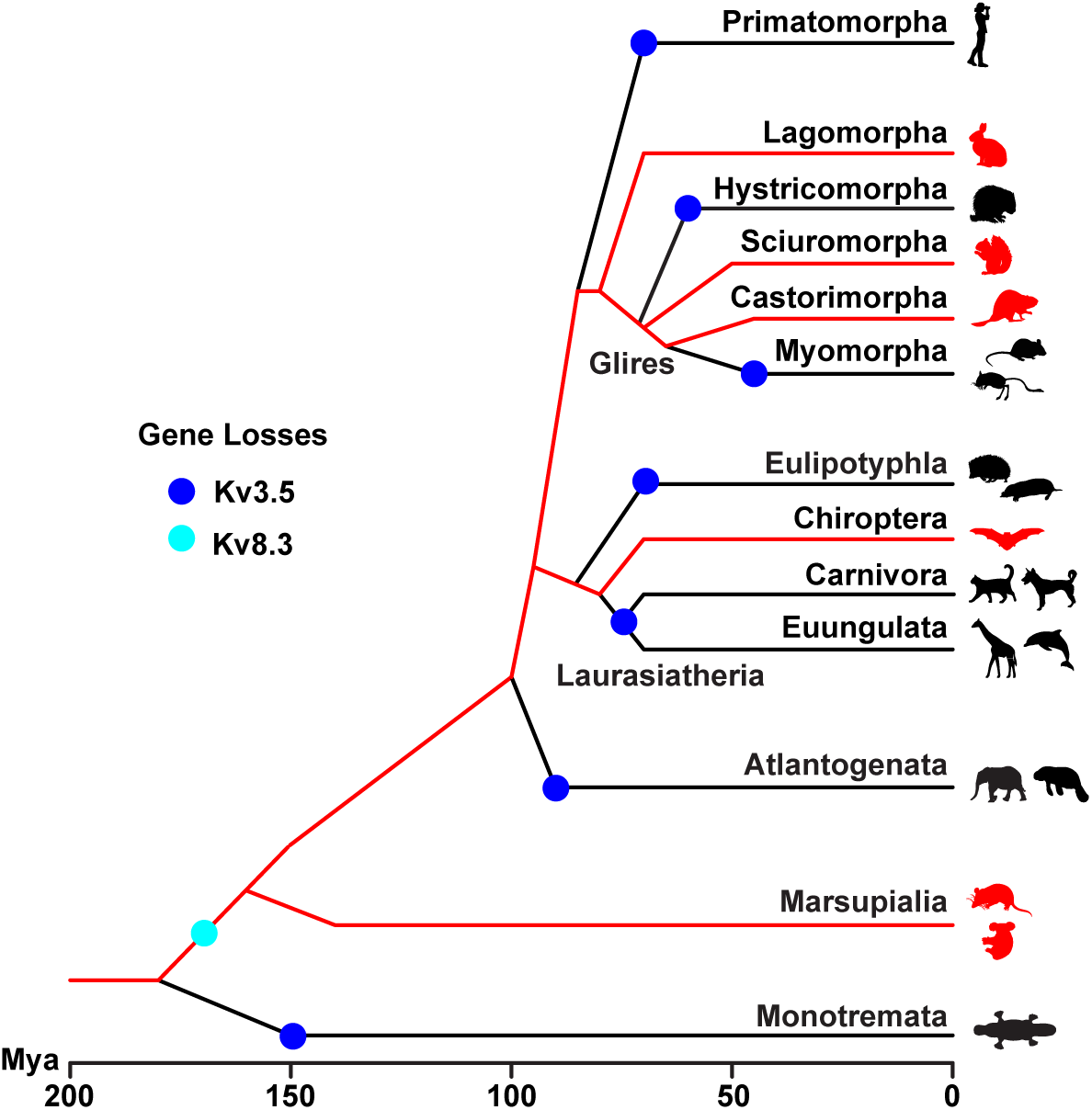
Loss of Kv3.5 and Kv8.3 in mammals. Schematic phylogeny of mammals based on current views of phylogenetic relationships and approximate divergence times in millions of years ago (Mya) from TimeTree 5 (Kumar et al., 2022). Mammalian clades are expanded to the extent needed to visualize major losses of Kv3.5. Lineages with Kv3.5 are highlighted in red and Kv3.5 losses are marked with 7 blue dots. The cyan dot marks the single loss of Kv8.3 after divergence of monotremes. Mammal icons at branch tips were obtained from the PhyloPic.org repository and are open-source images.

Kv3.3 and Kv3.4 inactivate via an N-type mechanism (Ruppersberg et al., 1991a; Fernandez et al., 2003) and we looked at the conservation of their N-terminal inactivation ball sequences to gain insights into the functional evolution of vertebrate Kv3s. The vertebrate Kv3 subfamily inactivation ball has cysteine and serine residues that render inactivation sensitive to oxidation and phosphorylation, respectively (Ruppersberg et al., 1991b; Covarrubias et al., 1994; Antz et al., 1997; Antz et al., 1999; Yang et al., 2019). Fast inactivation requires reducing conditions and an unphosphorylated ball. An alignment of Kv3 inactivation balls from representative vertebrate sequences is shown in Fig. 11C. We found conserved inactivation ball sequences with most or all modulation residues in all Kv3.3 and Kv3.4 orthologs as well as both clades of cyclostome Kv3s. This suggests the ancestral vertebrate Kv3 channel had N-type inactivation that was very likely already modulated post-translational modifications. Further evidence for ancestral inactivation comes from Kv3.1 orthologs in cartilaginous fish, the oldest extant gnathostome lineage, which also appear to have the inactivation ball (Fig. 11C, arrow). Assuming inactivation was indeed present in the ancestral vertebrate Kv3, then it was likely lost independently at different times in the Kv3.1 (late) and Kv3.2 (early) lineages. No invertebrate Kv3s had a similar inactivation ball, so rapidly inactivating Kv3 channels could be a vertebrate-specific evolutionary innovation.

The Kv3 subfamily has duplications in all the protostome lineages we examined, but it has not been duplicated in deuterostome invertebrates and has been lost in tunicates (Table 1). Most of the protostome duplications are limited to specific lineages; there are separate duplications in arthropods, nematodes and lophotrochozoans, for example. The Drosophila Kv3 channels Shaw and Shaw2 (Butler et al., 1989; Covarrubias and Rubin, 1993) have orthologs in insects and chelicerates, suggesting they originated from a duplication in the arthropod ancestor. The Kv3 subfamily is extensively expanded in cnidarians where the starlet sea anemone *Nematostella vectensis* has 12 Kv3s with evidence for 8 Kv3s in the cnidarian common ancestor (Lara et al., 2023). Teleosts have the most Kv3s in vertebrates (7 for zebrafish) due to 3R duplications of Kv3.1 and Kv3.3.

We expressed a Kv3.5 ortholog from vampire bat (Drotu-Kv3.5) in Xenopus oocytes to examine its functional properties. The most notable difference from Kv3.1-3.4 is that Drotu-Kv3.5 is a silent subunit that requires heteromeric assembly: it does not form functional channels on its own, but it is able to form functional heteromers when co-expressed with Drotu-Kv3.1 (Fig. 13A,B). Kv3.5 did not radically change the biophysical properties of Kv3.1: the heteromeric channel remains a high threshold delayed rectifier. But significant current size changes (Fig. 13B), a significant increase in GV slope (Fig. 13C, Table 2) and significant slowing of both activation and deactivation (Fig. 13D,E) provide strong evidence for functional heteromer formation between Drotu-Kv3.5 and Drotu-Kv3.1. Kv3.5 represents just the second independent evolution of silent subunits identified in the vertebrate Shaker family.

**Figure 13.**
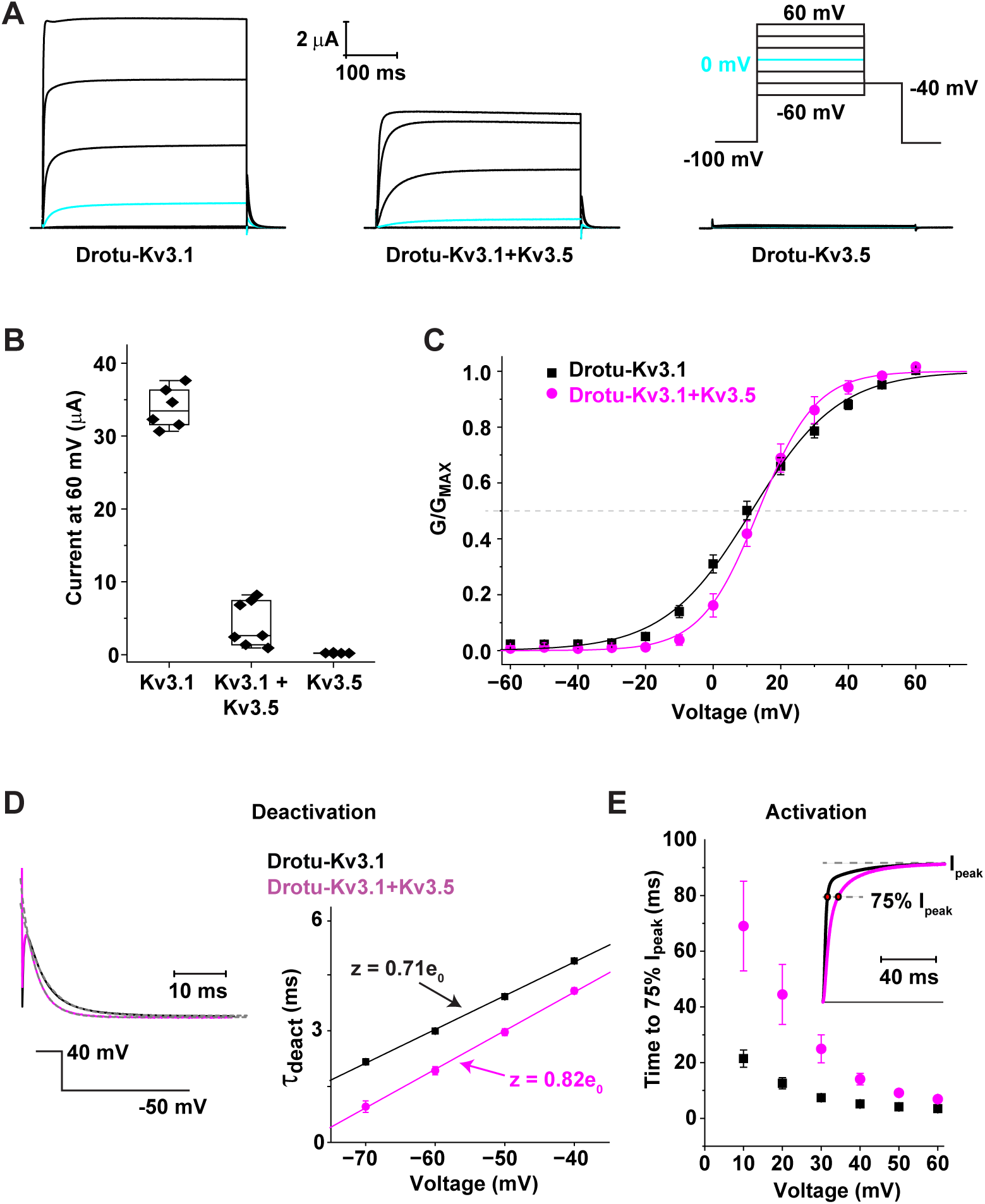
Functional characterization of Kv3.1 and Kv3.5 from vampire bat. (A) Example current traces recorded from oocytes expressing Drotu-Kv3.1 and Drotu-Kv3.5 alone and in combination. Scale bars are given for current amplitude and time and a schematic voltage protocol is shown at the upper right. (B) Box plots of peak current size at 60 mV for Drotu-Kv3.1, Drotu Kv3.1:Kv3.5 and Drotu-Kv3.5. Boxes show median and middle quartiles, whiskers mark the data range and data points from individual oocytes are depicted with black diamonds. Heteromeric Kv3.1:Kv3.5 currents were significantly smaller than Kv3.1 homomers (t-test, *p* < 0.0001) and Kv3.5 did not form functional channels. (C) Normalized GV curves for Kv3.1 homomers and Kv3.1/Kv3.5 heteromers measured from isochronal tail currents. Data points are mean ± S.D. and curves show Boltzmann fits using the average V_50_ and slope values determined from individual oocytes (Table 2). The GV slope was significantly changed in the Kv3.1:Kv3.5 heteromer (*p* < 0.05, t-test, n=6). (D) Deactivation time constants for Kv3.1 (black) and Kv3.1:Kv3.5 (magenta) measured from single exponential fits of tail currents recorded at −50 mV after test steps to 40 mV. Normalized example traces with overlaid exponential fits are shown in the left panel and a semi-log plot of time constant vs. voltage is shown in the right panel with exponential fits used to estimate the charge associated with deactivaton (z). (E) Plot of 75% activation time vs. voltage for Kv3.1 and Kv3.1:Kv3.5 with an inset to show normalized example traces. All data points in D and E were significantly different between the two channels except for deactivation at −40 mV (t-test, *p* < 0.05, n = 6-9).

### Shaker Kv4 Subfamily

The Kv4 subfamily BI and ML phylogenies showed consensus support for pan-gnathostome Kv4.1-3 clades in a (Kv4.2/Kv4.3)(Kv4.1) arrangement. Fig. 14 shows the BI phylogeny and the alignment and trees are provided as Data S34-36. Cyclostomes have a single Kv4 channel that pairs with the gnathostome Kv4.2 clade. Our interpretation is that the post-1R vertebrate ancestor still had a single Kv4 gene which was copied in the gnathostome 2R duplication and duplicated again independently in the stem gnathostome lineage. Kv4.1 was later lost in birds, and Kv4.2 and Kv4.3 were likely duplicated again in the teleost 3R genome duplication to produce 5 teleost Kv4 paralogs. The Kv4 subfamily has duplications in three invertebrate lineages: cnidarians, chelicerates and cephalochordates. The cnidarian duplications include silent subunits that require heteromeric assembly (Jegla and Salkoff, 1997; Li et al., 2015b; Lara et al., 2023). Kv4 duplications in cephalochordates are extensive and some of the genes are highly divergent. We therefore included only a subset of cephalochordate Kv4s in our analyses to preserve alignment length.

**Figure 14.**
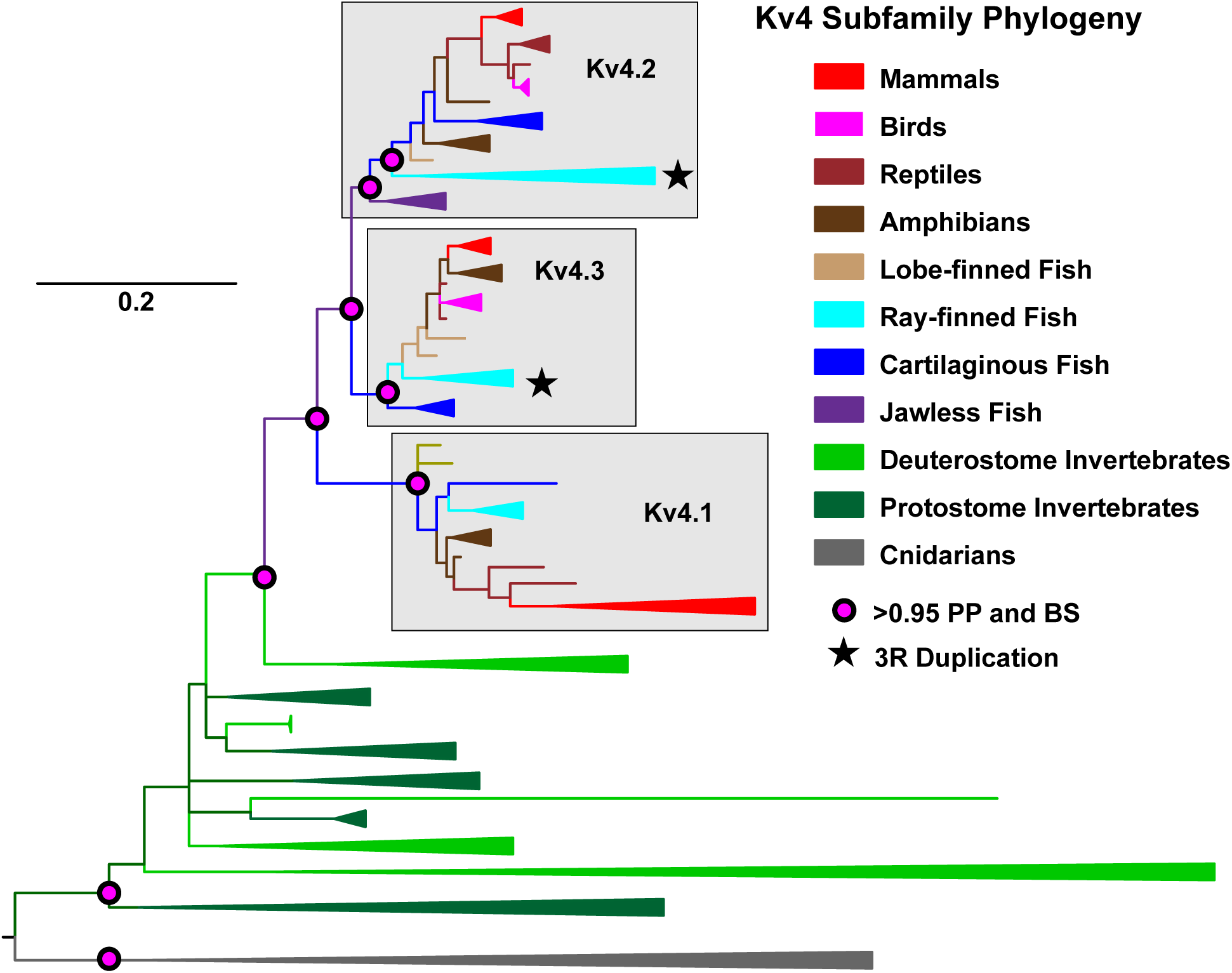
Phylogenetic analysis of the Kv4 subfamily. Bayesian inference phylogeny of the Kv4 subfamily phylogeny containing 158 channels in a 418-site alignment. It is rooted between cnidarians and bilaterians for display. Clades are colored according to the legend, collapsed clade width does not reflect sequence number and a scale bar is provided for substitutions/site. Pan-gnathostome ortholog clades are shaded and labeled Kv4.1-4.3. Support for important nodes is indicated with colored circles for this phylogeny and a ML version run on the same alignment. Stars mark putative teleost 3R duplications in the Kv4.2 and Kv4.3 clades.

## DISCUSSION

We found a total of 45 Kv family channels with broad distribution in gnathostome vertebrates, including 6 from the KCNQ family, 9 from the EAG family and 30 from the Shaker family. The phylogenetic distribution of these channels is summarized in Fig. 15A. 43 of these channels were present in the gnathostome common ancestor; only Kv12.2 and Kv3.5 arose from later duplications. Kv12.2 is the youngest human Kv channel with an origin in a common ancestor of lobe-finned fish and tetrapods. The Kv2 subfamily silent subunit Kv5.1 is the oldest human Kv channel it appeared in the chordate ancestor of vertebrates and tunicates and has never been duplicated in the human lineage. All other human Kv channels are the product of duplications that occurred within the vertebrate lineage and do not have 1:1 orthologs in any invertebrate.

**Figure 15.**
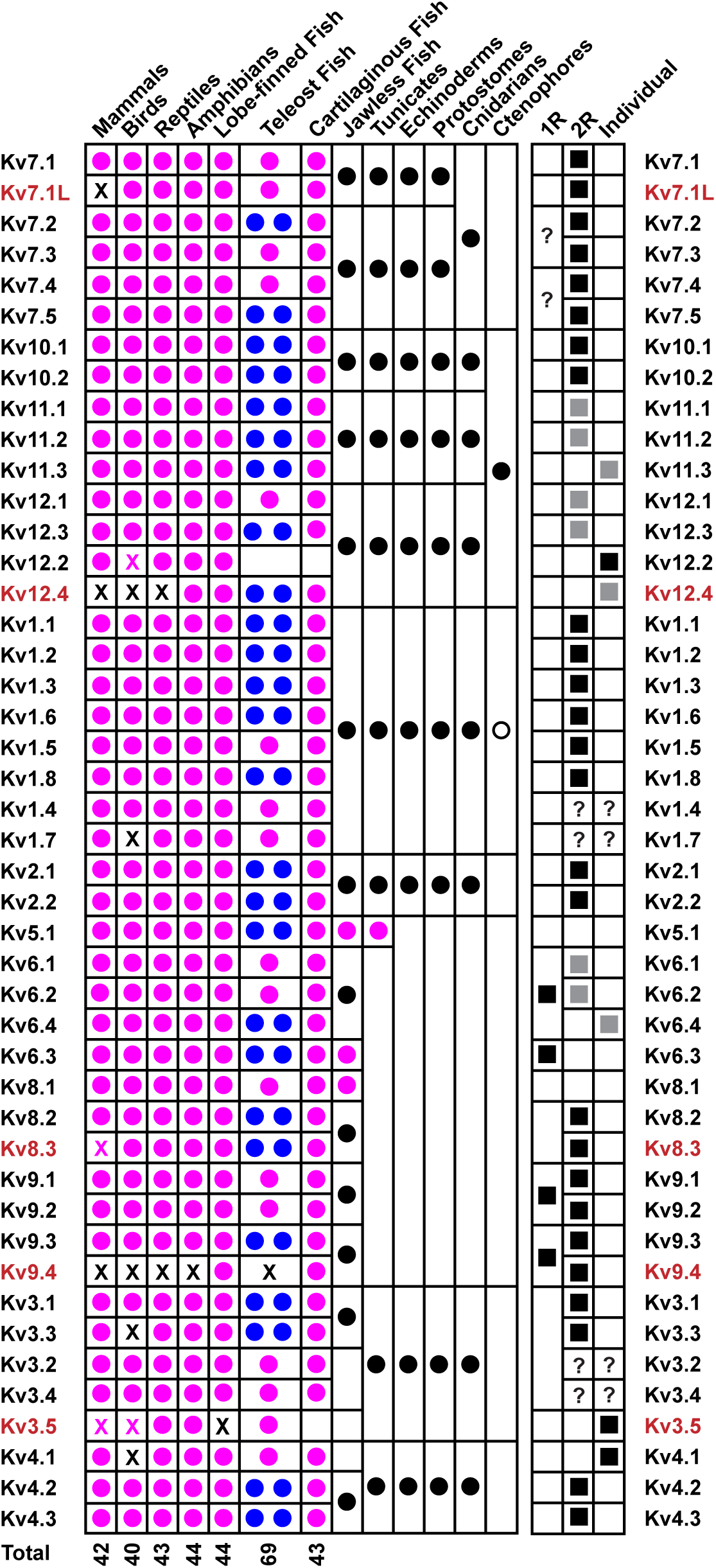
Summary of the phylogenetic distribution of vertebrate Kv orthologs. Rows represent 45 vertebrate Kv channel genes (labeled at left and right) that were present at some point in the human evolutionary lineage. Columns in the left block show phylogenetic distribution. Column in the right block indicate duplication history. Magenta dots indicate the presence of 1:1 orthologs and Xs mark gene losses the indicated gnathostome clade (black, complete loss; magenta, partial loss). Double blue dots are used to mark genes with evidence for 3R duplications in teleost fish. Total number of Kv genes predicted for the ancestor of each gnathostome clade is indicated at the bottom. Black dots are used to indicate presence of channels that are not in 1:1 orthology relationships with gnathostome channels but are specifically related to the gnathostome channels bracketed by their box borders. For example, jawless fish have KCNQ1 and KCNQ2 subfamily channels but not 1:1 orthologs of 6 gnathostome channels. Ctenophores Shakers (open circle) share more characteristics with the Kv1 subfamily than the Kv2-4 subfamilies, but are not true Kv1s (Li et al., 2015b; Jegla et al., 2024; Simonson et al., 2024; Simonson et al., 2025). Black squares in the right block are used to indicate a putative role for 1R, 2R or independent duplications in the evolutionary history of the 45 Kv orthologs. Question marks are used when the role is highly uncertain and gray squares indicate uncertainty about which of the genes are the 2R ohnologs and which were independently duplicated.

We assume Kv5.1 acquired the silent phenotype no later than in the stem vertebrate lineage before the diversification of the major Kv2 subfamily silent subunit clades because our phylogenetic analyses support a model in Kv6, Kv8 and Kv9 channels are duplicated from an ancestral Kv5 channel. Functional characterization of tunicate Kv5.1 orthologs could reveal whether the silent phenotype predates vertebrates. Until our characterization of Kv3.5 here, these Kv2 subfamily silent subunits were the only known vertebrate Shaker family silent subunits.

However, silent subunit evolution happened multiple times in cnidarians, and Shaker family silent subunits are also present in ctenophores and choanoflagellates (Lara et al., 2023; Jegla et al., 2024; Simonson et al., 2025). Interestingly, the original assembly partner of the vertebrate Kv2 subfamily silent subunits must have been the single ancestral Kv2 subfamily α-subunit our phylogenies predict rather than Kv2.1 and Kv2.2 which arose much later from a gnathostome-specific duplication. The extended period of co-evolution between this ancestral Kv2 α-subunit and multiple clades of silent subunits may explain why both Kv2.1 and Kv2.2 can both assemble efficiently with silent subunits.

A summary of the role of the stem vertebrate 1R genome duplication and the gnathostome-specific 2R genome duplication in building the human Kv channel set is shown in Fig. 15B. We found little evidence for contribution of the 1R genome duplication to the human Kv set. Our phylogenetic analyses suggest the vertebrate common ancestor had 14 Kv channels prior to 1R: one each of Kv1-6 and Kv9-12, two Kv7s and two Kv8s. If we assume that all these 14 genes were initially duplicated in 1R, then we could expect a set of up to 28 Kv channels shared between cyclostomes and gnathostomes. Instead, we find evidence for unambiguous 1R duplications only for the Kv6 and Kv9 lineages (Fig. 8), bringing the stem vertebrate Kv channel total to just 16 genes. The presence of (AB)(CD) topologies for the gnathostome Kv7.2-7.5 and Kv3.1-3.4 clades suggests that these expansions might also include 1R duplications despite that absence of cyclostome sequences paired with each AB and CD clade (Fig. 2,11). If so, that would bring the post-1R Kv channel total to 18. The retention rate for additional 1R ohnologs was therefore probably only 14-29% (2−4/14). It is alternatively possible that the Kv7.2-7.5 and Kv3.1-3.4 clades were each expanded by an independent duplication between 1R and 2R to reach 18 genes.

We can infer the gnathostome Kv channel set had further increased to as many as 21 genes prior to the 2R gnathostome genome duplication with the addition of the Kv1 subfamily 3-gene array. The pre-2R Kv1 subfamily would have included a total 4 genes because we infer it must also have had a singleton to eventually give rise to Kv1.4 and Kv1.7. We would therefore predict a potential for up to 42 gnathostome 2R ohnologs. We find good evidence in the phylogenies for 32 2R ohnologs based on statistically supported pairs of pan-gnathostome Kv ortholog groups in the phylogenies, and there are 3 unambiguous cases where 2R ohnologs were lost: Kv5.1, Kv8.1 and Kv6.3. Kv1.4/Kv1.7 could represent four 2R ohnologs based on phylogenetic distribution, but support is weaker because they don’t form pairs with consensus statistical support. The pair could instead have originated from a post 2R individual duplication. The Kv3.2 and Kv3.4 origins are even less clear, but their presence requires an individual duplication either before or after 2R, and then another duplication via 2R or 2^nd^ post-2R individual duplication. We include a Kv3.2/3.4 ancestor in the 21 gene pre-2R Kv count, so if this ancestor was instead generated by a post-2R duplication, it would reduce the pre-2R Kv channel set to 20 genes with a potential for 40 2R ohnologs. Therefore, if we assume one extreme that Kv1.4, Kv1.7, Kv3.2 and Kv3.4 all arose from post 2R duplications, then the retention rate for 2R additions to the Kv set 16/20 or 80%. If we instead assume the other extreme - that Kv1.4/Kv1.7 and Kv3.2/Kv3.4 are 2R ohnolog pairs - then retention was 18/21 or 86%. Regardless of the precise scenario, the evidence is clear that 2R Kv ohnologs were retained at a much higher rate than 1R Kv ohnologs. Without those 1R ohnolog losses, the human Kv set might have been much larger.

Additional post-2R gene duplications in the stem gnathostome added new Kv4, Kv6, Kv11 and Kv12 ortholog groups for a total of 43 pan-gnathostome Kv channels. The further addition of Kv3.5 raised the total Kv channel complement to 44 in Osteichthyes before the divergence of tetrapods and ray-finned fish. The addition of Kv12.2 in lobe-finned fish raised the total to 45 Kv channels in the Sarcopterygii prior to the emergence of tetrapods. We found evidence for up to 69 Kv genes in the teleost ancestor after the 3R genome duplication based on the presence of co-ortholog pairs for 25/44 (57%) of the ancestral ray-finned fish Kv genes produced by the 3R genome duplication. This number is a very rough estimate because we did not follow up with a deeper coverage phylogenetic analysis of teleost fish species that would be necessary to more confidently sort 3R duplications from isolated lineage-specific duplications.

Humans lost 5/45 Kv channels present in the stem tetrapod, with three of the losses occurring within the mammal lineage: Kv7.1L is absent in all mammals but present in birds and reptiles, Kv8.3 is present in monotremes but lost in marsupials and placental mammals, and Kv3.5 was lost multiple times in mammalian evolution but is still present in some species today. Mice lost the same 5 channels as humans, but Kv3.5 losses in the mouse and human lineages stems are independent. The mammalian common ancestor therefore had 42/45 tetrapod Kv channels, but living mammals have only 40-41 Kvs due to ongoing losses. Birds also had significant Kv losses (Fig. 15A) and we predict the bird ancestor had just 40 Kv channels. Kv1.7, Kv3.3, Kv4.1, Kv9.4 and Kv12.4 are entirely absent in birds. The Kv9.4 and Kv12.4 losses occurred prior to divergence from reptiles, but the other three losses are specific to birds. Most extant bird species have also lost Kv12.2 and Kv3.5 and thus have only 38 Kv channels. The stability of the ancestral gnathostome Kv channel set is nevertheless remarkable with 36/43 channels conserved in all major gnathostome lineages with very few new additions outside teleost fish.

Biophysical characterization of the five Kv channels humans lost suggests these gene losses did not punch any major fundamental holes in the functional breadth of the human Kv set. Onat-Kv8.3 and Lchal-Kv9.4 are highly similar to remaining human Kv8s and Kv9s in terms of gating to the point that they could probably be assigned to the correct silent subunit clades if their molecular identities were unknown. One of the more interesting conclusions from this is that the functional properties of the Kv2 silent subunit clades differentiated very early in vertebrate evolution before the 1R and 2R genome duplications. Further evidence for this comes from conservation of comparatively large hyperpolarizing GV and SSI shifts observed across the Kv6 clade (Post et al., 1996; Kramer et al., 1998; Zhu et al., 1999; Bocksteins et al., 2012; Pisupati et al., 2018). Drotu-Kv3.5 is clearly unique compared to other human Kv3s in that it is a silent subunit. However, the functional consequences of co-assembly between Drotu-Kv3.1 and Drotu-Kv3.5 are relatively unremarkable. The heteromeric current is still a high threshold delayed rectifier and the changes in kinetics are modest. While we did not have a Kv7.1 clone for direct comparison, the Hharp-Kv7.1L current is also remarkably similar to published descriptions of Kv7.1, with the only major difference being the polarity of the GV shift induced by Hsapi-KCNE1. This might not be a major loss in terms of biophysical diversity of KCNQ1 subfamily channels, however, since Kv7.1 activation can be shifted in a hyperpolarizing direction by KCNE2 and KCNE3 (Schroeder et al., 2000; Tinel et al., 2000; Barro-Soria et al., 2015).

The most biophysically unique of the lost channels is Amexi-Kv12.4 which has unusually rapid deactivation for a vertebrate Kv12 channel. Humans nevertheless have multiple kinetically diverse Kv channels that operate over a similar voltage range including Kv12.2, Eag and Erg subfamily channels, and even KCNQ family channels. Furthermore, the teleost Kv12.4 ortholog zElk and human Kv12.1 do share open state stabilization upon prolonged depolarization that can slow of deactivation (Li et al., 2015a; Dai and Zagotta, 2017). Stabilization requires PIP2 and depends on eag domain/CNBHD interactions, so the kinetics and voltage-activation range of these Kv12 channels in native cells might therefore be dynamically regulated. We suggest PIP2-dependent open state stabilization could be an ancestral property of vertebrate Elk channels because the last common ancestor of Kv12.1 and Kv12.4 is the single ancestral vertebrate Kv12 channel predicted from phylogenetic analyses. However, open state stabilization remains to be studied in Kv12.2 or Kv12.3.

The human Kv set remains remarkably diverse on a biophysical level even without the five lost channels, and it is therefore hard to argue that we lost them because we don’t need as much Kv current diversity as our ancestors. Numerous functional studies of diverse invertebrate and vertebrate Kv demonstrate the basic gating phenotypes of our Kv1-12 channel clades were already set prior to the gnathostome-specific duplications that expanded our Kv channel set.

Thus, human Kv paralogs are mostly modest functional variations on ancestral gating phenotypes, and one can argue that there is therefore substantial gating redundancy remaining in the human Kv channel set. There are few if any truly novel gating phenotypes that arose in Kv channels as within the gnathostome lineage despite the addition of so many ohnologs.

The more relevant question then becomes why did we keep so many of the channels produced by Ohno’s genome duplications? The answer most likely lies in the partitioning of function and subfunctionalization that genome duplications enable (Wolfe, 2001). Loss-of-function mutations that restrict expression pattern of individual paralogs are the canonical example of partitioning: if a channel type is needed in all neurons, these partitioning mutations would not be tolerated prior to duplication. But after duplication, expression patterns can degenerate for individual paralogs so long as the full paralog set still covers all neurons.

Partitioning of expression pattern can force retention of multiple paralogs as it extends across the set. There are numerous descriptions of unique expression patterns for human and mouse Kv paralogs in the literature that might be examples of partitioning. For example, each of the 8 mammalian Kv10-12 paralogs in the EAG family has a unique neuronal expression pattern (Shi et al., 1997; Shi et al., 1998; Saganich et al., 1999; Saganich et al., 2001; Zou et al., 2003).

A case of subfunctionalization in Kvs, where multiple paralogs are needed to produce sufficient functional protein, could be the co-assembly of Kv7.2 and Kv7.3 to produce the classic neuronal m-current (Wang et al., 1998). Co-assembly of Kv7.2 and Kv7.3 is more efficient than homomeric assembly, and it is required to produce sufficient m-current in neurons because further loss-of-function in either subunit leads to epilepsy in humans (Charlier et al., 1998; Singh et al., 1998). The single Kv7.2/7.3 ancestor prior to the 2R genome duplication must have had efficient homomeric assembly, so the reduced homomeric assembly efficiency in Kv7.2 and Kv7.3 probably stems from loss-of-function mutations that were no longer deleterious post-duplication. We suggest loss-of-function mutations that led to partitioning and subfunctionalization were probably a key driver behind the long-term evolutionary stability of the gnathostome Kv channel set. Partitioning of expression pattern would have in turn provided more opportunities for functional tuning of individual vertebrate Kv paralogs to specific cell types, which we may now observe as modest gating variations between paralogs. We suggest that evolutionary changes in the expression patterns or levels of Kv3.5, Kv7.1L, Kv8.3, Kv9.4 and Kv12.4 could therefore be relevant to understanding why these specific channels were lost from the human lineage, and it would be interesting to explore these in future studies.

## Data Availability Statement

Data shown in Fig. 2-15 are available in the article and online supplemental materials. Electrophysiology recording files can be obtained from the corresponding author upon reasonable request.

## Acknowledgments

This work was made possible by a diverse research team and the unique perspectives each member brought to the table. We are greater than the sum or our parts. Research was supported by the Huck Institutes of the Life Sciences and the Eberly College of Science at Penn State University.

