## Supplementary material for "Tracing the origins of human voltage-gated K^+^ channels: where they come from and what we lost along the way": Table S1

| Table S1. Phylogeny run parameters | | | | | | | | |
|---|---|---|---|---|---|---|---|---|
| | Alignment | | Bayesian Inference | | | | | Maximum Likelihood |
| Phylogeny | Taxa | Sites | Model | PSFR[1] | Min. ESS[2] | Ave. ESS | S.D.[3] | Model |
| KCNQ | 244 | 398 | JTT[4] | 1.00 | 2063 | 2379 | 0.009 | LG[5]+R7 |
| KCNQ2 (vert) | 128 | 530 | JTT | 1.00 | 1124 | 1218 | 0.012 | Q.mammal[6]+R5 |
| EAG | 446 | 596 | JTT | 1.00 | 895 | 1112 | 0.031 | LG+F+I+R9 |
| Kv1 Full | 356 | 330 | JTT | 1.00 | 4385 | 4985 | 0.017 | LG+R8 |
| Kv1 (vert) | 271 | 389 | JTT | 1.00 | 6360 | 7091 | 0.011 | Q.mammal+I+R7 |
| Kv2 Full | 509 | 393 | JTT | 1.00 | 1732 | 2193 | 0.056 | LG+F+R10 |
| Kv2.1/2.2 | 74 | 681 | JTT | 1.00 | 1054 | 1109 | 0.012 | Q.mammal+R5 |
| Kv3 | 235 | 373 | JTT | 1.00 | 3113 | 3219 | 0.017 | LG+I+R6 |
| Kv3 (Vert) | 158 | 484 | JTT | 1.00 | 1546 | 2446 | 0.014 | Q.mammal+I+R6 |
| Kv4 | 167 | 418 | WAG[7] | 1.00 | 2750 | 2850 | 0.014 | LG+I+R6 |

[1]Potential Scale Reduction Factor, 1.00 indicates convergence of runs

[2]Effective Sample Size, >200 indicates good chain mixing during the run

[3]Standard Deviation of the Split Frequencies, measures agreement between chains. Below 0.05 is good for large phylogenies with groups of closely related orthologs that are difficult to differentiate. The higher value for the full Kv2 subfamily alignment reflects the inability to resolve order in the large vertebrate Kv2.1 and Kv2.2 clades.

[4] (Jones et al., 1992)

[5] (Le and Gascuel, 2008)

[6] (Minh et al., 2021)

[7] (Whelan and Goldman, 2001)

### TABLE REFERENCES


Jones, D.T., W.R. Taylor, and J.M. Thornton. 1992. The rapid generation of mutation data matrices from protein sequences. *Comput Appl Biosci*. 8:275-282.

Le, S.Q., and O. Gascuel. 2008. An improved general amino acid replacement matrix. *Molecular biology and evolution*. 25:1307-1320.

Minh, B.Q., C.C. Dang, L.S. Vinh, and R. Lanfear. 2021. QMaker: Fast and Accurate Method to Estimate Empirical Models of Protein Evolution. *Systematic biology*. 70:1046-1060.

Whelan, S., and N. Goldman. 2001. A general empirical model of protein evolution derived from multiple protein families using a maximum-likelihood approach. *Molecular biology and evolution*. 18:691-699.
